# Areas of global importance for terrestrial biodiversity, carbon, and water

**DOI:** 10.1101/2020.04.16.021444

**Authors:** Martin Jung, Andy Arnell, Xavier de Lamo, Shaenandhoa García-Rangel, Matthew Lewis, Jennifer Mark, Cory Merow, Lera Miles, Ian Ondo, Samuel Pironon, Corinna Ravilious, Malin Rivers, Dmitry Schepashenko, Oliver Tallowin, Arnout van Soesbergen, Rafaël Govaerts, Bradley L. Boyle, Brian J. Enquist, Xiao Feng, Rachael V. Gallagher, Brian Maitner, Shai Meiri, Mark Mulligan, Gali Ofer, Jeffrey O. Hanson, Walter Jetz, Moreno Di Marco, Jennifer McGowan, D. Scott Rinnan, Jeffrey D. Sachs, Myroslava Lesiv, Vanessa Adams, Samuel C. Andrew, Joseph R. Burger, Lee Hannah, Pablo A. Marquet, James K. McCarthy, Naia Morueta-Holme, Erica A. Newman, Daniel S. Park, Patrick R. Roehrdanz, Jens-Christian Svenning, Cyrille Violle, Jan J. Wieringa, Graham Wynne, Steffen Fritz, Bernardo B.N. Strassburg, Michael Obersteiner, Valerie Kapos, Neil Burgess, Guido Schmidt-Traub, Piero Visconti

## Abstract

To meet the ambitious objectives of biodiversity and climate conventions, countries and the international community require clarity on how these objectives can be operationalized spatially, and multiple targets be pursued concurrently^1^. To support governments and political conventions, spatial guidance is needed to identify which areas should be managed for conservation to generate the greatest synergies between biodiversity and nature’s contribution to people (NCP). Here we present results from a joint optimization that maximizes improvements in species conservation status, carbon retention and water provisioning and rank terrestrial conservation priorities globally. We found that, selecting the top-ranked 30% (respectively 50%) of areas would conserve 62.4% (86.8%) of the estimated total carbon stock and 67.8% (90.7%) of all clean water provisioning, in addition to improving the conservation status for 69.7% (83.8%) of all species considered. If priority was given to biodiversity only, managing 30% of optimally located land area for conservation may be sufficient to improve the conservation status of 86.3% of plant and vertebrate species on Earth. Our results provide a global baseline on where land could be managed for conservation. We discuss how such a spatial prioritisation framework can support the implementation of the biodiversity and climate conventions.

## Introduction

Biodiversity and nature’s contributions to people (NCP) are in peril, requiring an increasing level of ambition to avert further decline^1^. Existing global biodiversity conservation targets are unlikely to be met by the end of 2020^2^. Similarly, the world is falling short of mobilizing the full climate mitigation potential of nature-based climate solutions, estimated at around a third of mitigation effort under the Paris Agreement^3^. A new global biodiversity framework is scheduled to be adopted by the Convention on Biological Diversity (CBD) in Kunming, China, in October 2020^4^, and there are growing calls to integrate nature-based solutions into climate strategies^5^.

Targets for site-based conservation actions, hereafter area-based conservation targets, will likely remain important for the new global biodiversity framework^4^. Several calls have been made for such targets, including suggestions that at least 30% of land and oceans be protected for conservation and an additional 20% for climate mitigation^6^ and that the value of areas of global importance for conservation is maintained or restored^7^. The Sustainable Development Goals (SDGs), the United Nations Framework Convention on Climate Change (UNFCCC) and the CBD emphasize that habitat conservation and restoration should contribute simultaneously to biodiversity conservation and climate change mitigation^4^. Recent analyses of conservation priorities for biodiversity and carbon have spatially overlaid areas of importance for both assets, effectively treating the two goals as to be pursued separately (e.g.^6,9^). However, multi-criteria spatial optimization approaches applied to conservation and restoration prioritisation have shown that carbon sequestration could be doubled, and the number of extinctions prevented tripled, if priority areas were jointly identified rather than independently^10,11^. Yet, no comparable optimization analyses exist at a global scale.

A number of recent studies have attempted to map spatial conservation priorities on land^12^, relying on spatial conservation prioritisation (SCP) methods^13–16,17^. However, these approaches are limited, in that: they (*i*) are limited by geographic extent^22^ or focus on only a subset of global biodiversity, notably ignoring either reptiles or plant species, which show considerable variation in areas of importance compared to other taxa ^18,19^; (*ii*) focus on species representation only, rather than reducing extinction risk, as per international biodiversity targets, and often ignore other dimensions of biodiversity, e.g. evolutionary distinctiveness^20,21^; (*iii*) do not investigate the extent to which synergies between biodiversity and NCPs, such as carbon sequestration or clean water provisioning^22^, can be maximised^21^; and (*iv*) they use a-priori defined, and subjective measures of importance, such as intactness^8,17^, or area-based conservation targets, such as 30% or 50% of the Earth^6,24^ instead of objectively delineating the relative importance of biodiversity and NCPs across the whole world irrespective of such constraints.

The aim of this study is to identify the most important areas for biodiversity - here focussing on species conservation - as well as NCPs including carbon storage and water provisioning, to be managed for conservation globally. We define managing an area for conservation as any site-based action that is appropriate for the local context (considering pressures, tenure, land-use, etc.), and that is commensurate with retaining or restoring the desirable assets (e.g. species, habitat types, soil or biomass carbon, clean water). This management may sometimes require legal protection to be effective, but not necessarily in the form of protected areas.

We obtained fine-scale distribution maps for the world’s terrestrial vertebrates as well as the largest sample of plant distribution data ever considered in global species-level analysis, ~41% of all accepted species names in this group. As NCPs we use the latest global spatial data on above- and below-ground biomass carbon, and vulnerable soil carbon, as well as the volume of potential clean water by river basin. We applied a multicriteria spatial optimization framework to investigate synergies between these assets and explore how priority ranks change depending on how much weight is given to either carbon sequestration, water provisioning or biodiversity, and examined whether priorities vary if species evolutionary distinctiveness and threat status are considered.

## Results

We found large potential synergies between managing land for biodiversity conservation, storing soil and biomass carbon, and maintaining clean water provisioning. Managing the top-ranked 10% of land, i.e. those areas with the highest priority, to achieve these objectives simultaneously (Fig. 1, SI Fig. 1), has the potential to improve the conservation status of 46.1% of all species considered, of which 51.1% are plant species, as well as conserve 27.1% of the total carbon and 24.1% of the potential clean water globally. Areas of biodiversity importance notably include mountain ranges of the world, large parts of Mediterranean biomes and South-East Asia (SI Fig. 2) and were overall mostly comparable to previous expert-based delineations of conservation hotspots^16^, while also highlighting additional areas of importance for biodiversity only, such as the West African Coast, Papua New-Guinea and East Australian Rainforest (SI Fig. 2). The Hudson Bay area, the Congo Basin and Papua New Guinea were among the top-ranked 10% areas for global carbon storage (SI Fig. 3a), while the Eastern United States of America, the Congo, European Russia and Eastern India were among the areas with the greatest importance for clean water provisioning (SI Fig. 3b). Overall, top-ranked areas of joint importance of biodiversity, carbon and water were spatially distributed across all continents, latitudes and biomes.

**Fig. 1:**
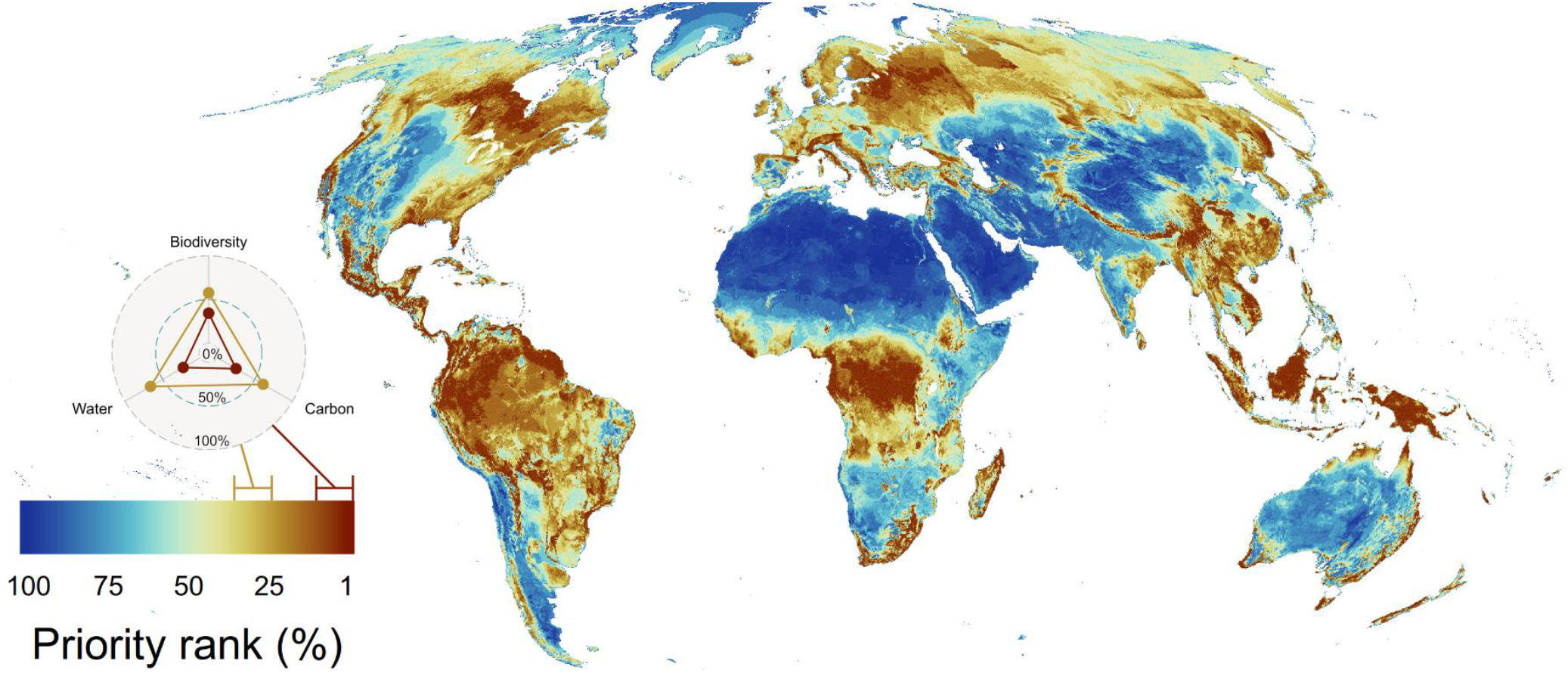
Global areas of importance for terrestrial biodiversity, carbon and water. All assets were jointly optimized with equal weighting given to each asset (central point in the series of segments in Fig. 2) and ranked by the most (1-10%) to least (90-100%) important areas to conserve globally. The triangle plot shows the extent to which protecting the top-ranked 10% and 30% of land (dark brown and yellow areas on the map) contributes to improving species conservation status, storing carbon and providing clean water. The map is at 10 km resolution in Mollweide projection. A map highlighting the uncertainty in priority ranks can be found in SI Fig 1.

Synergies and trade-offs depend on the relative importance given to conservation of terrestrial biodiversity, carbon storage and water provisioning (Fig. 2a). We explored an array of conservation scenarios each with a range of possible outcomes: at one extreme, priority is given to conserving biodiversity and carbon only, and with equal weight (Fig. 2b). At the other extreme are scenarios that prioritize conserving only biodiversity and water (Fig. 2c). Intermediate options include giving equal weighting to all three assets (Fig. 1). Similar to earlier assessments^9,26,27^, we found synergies between the conservation of biodiversity and carbon storage (Fig. 2b). However we also discovered similar synergies for biodiversity and water provisioning (Fig. 2c). Conserving the top-ranked 10% of land for biodiversity and carbon can only protect up to 23.6% of the global total carbon and 45.8% of all species (Fig 2a), while maintaining 17.8% of all global water provisioning as co-benefit (Fig. 2b). In contrast, conserving the top-ranked 10% of land for biodiversity and water only can protect 21.7% of water and 43.6% of all species (Fig 2a), while maintaining 18% as carbon co-benefit (Fig. 2c). The implications of assigning different relative preferences to conserving NCPs magnify with increasing amounts of land dedicated to conservation. For example, with 10% and 30% of land managed for conservation the range of carbon conserved is 18% to 23.6% and 49.2% to 63.1% respectively, and the range in water conserved is 17.8% to 21.7% and 51.8% to 66.4% (Fig. 2a). Our results suggest that there is ample scope for identifying co-benefits from conserving these three assets, if explicit targets for each are considered, areas of importance for each asset are identified through multi-criteria optimization, and the range of relative weights given to each asset is comprehensively explored.

**Fig. 2:**
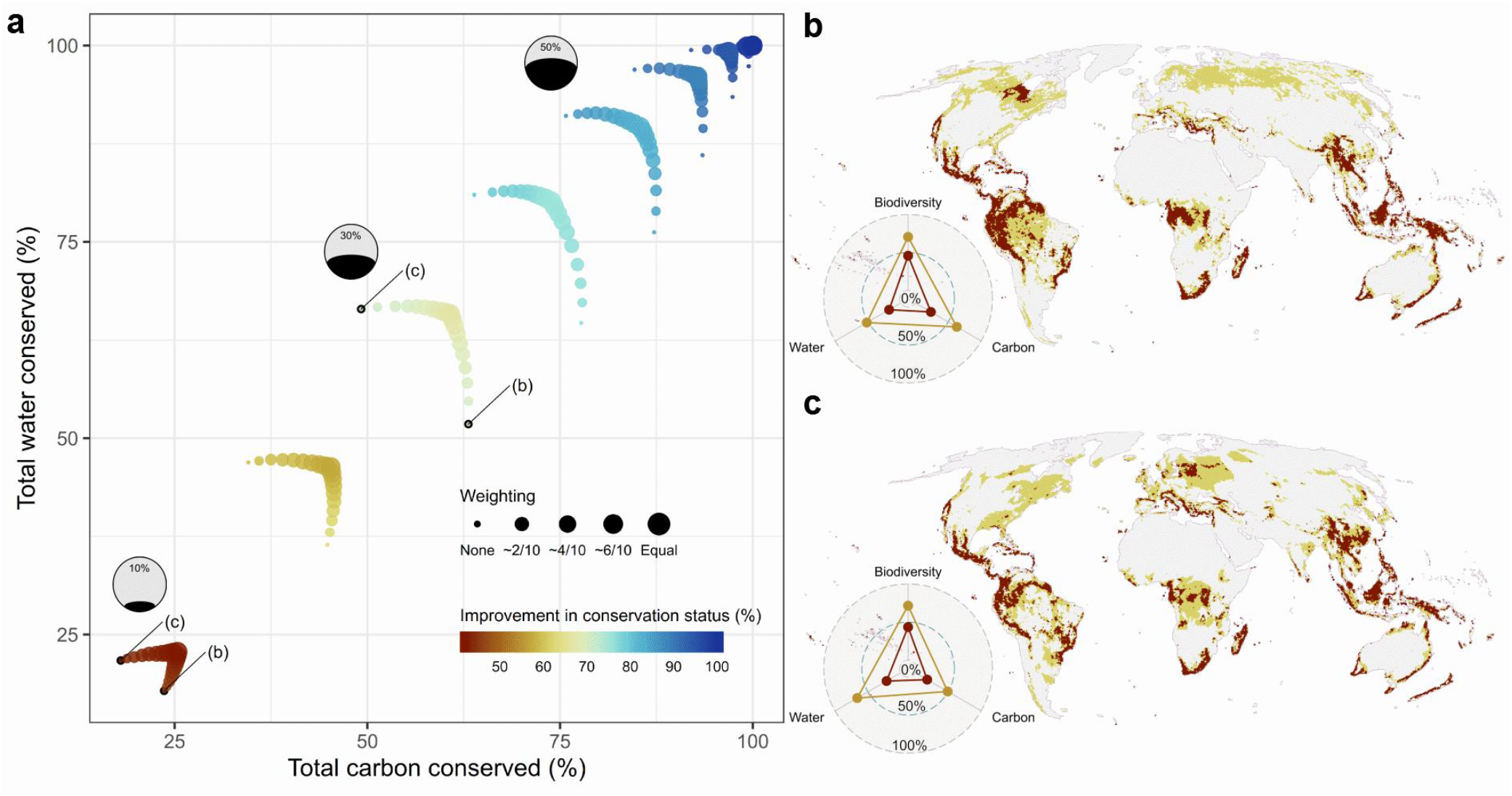
Implications of different relative weights given to carbon or water over improving species conservation status. (a) Each ‘boomerang-shaped’ segment of dots represents a series of conservation prioritisation scenarios with a common area budget (from 10% of land bottom left to 100% at top-right). Axes indicate the proportion of all carbon and water provisioning assets conserved, colours represent the proportion of species for which conservation status could be improved in a given conservation scenario, and the point size indicates the difference in weighting given to carbon or water relative to biodiversity, ranking from none to equal weighting. (b-c) Global areas of importance if 10% (dark-brown), or 30% (yellow), of land area is managed for conservation while preferring (b) carbon protection over water or (c) water protection over carbon.

The amount of land necessary to exclusively protect global biodiversity continues to be debated^15,28,29^ In our analysis we found that, in the absence of any socio-economic constraints and ignoring other NCPs (here water and carbon), at least ~67% of land needs to be managed for conservation globally, to improve the conservation status for terrestrial plants and vertebrates (Fig. 3a). This is robust to the number of species included in the analyses, provided that they are a representative subset (see methods), with the variation typically being ~0.1% around the mean accumulation curves (Fig. 3a).

**Fig. 3:**
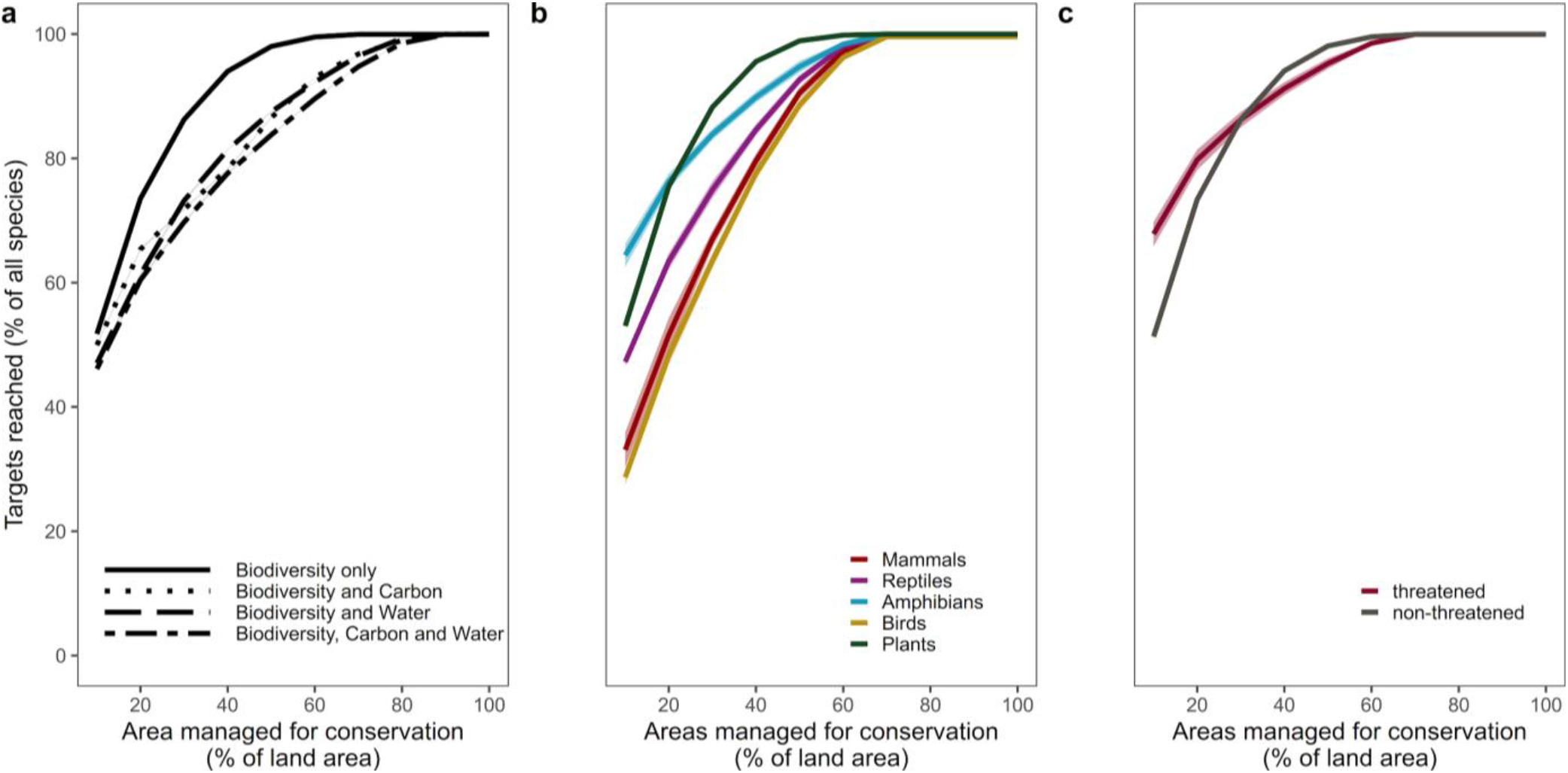
Accumulation curves showing how the number of species targets met increases with amount of land optimally allocated to conservation. Confidence bounds of accumulation curves indicate the uncertainty among representative sets and were generally found to be very small (~0.1%). This analysis ignores current protected areas and a version including those areas can be found in the SI Fig. 6. (a) Target accumulation curves for analysis variants including other assets; (b) for different taxonomic groups when optimizing biodiversity only to conservation; (c) for species classified by IUCN as threatened or not (see Methods) when optimizing for biodiversity only.

Optimally placing areas managed for conservation on 30% of the world’s land is already sufficient to conserve 86.3% of all species considered in this analysis (ignoring existing protected areas, socio-economic constraints and other NCPs). Currently protected areas (PAs) are potentially sufficient to achieve persistence targets for 16.3% of the species analysed (SI Fig. 5, SI Fig. 6). However, by building on the current PA estate to increase areas managed for biodiversity conservation up to 30% of land, the conservation status of an additional 60.8% of the species could be improved (for a total of 77.1% of the species analysed). Therefore, there is an efficiency gap of only ~9.2% between re-designing global conservation efforts and optimally building on existing efforts.

When jointly optimizing target achievement for biodiversity, carbon and water (Fig. 3a), we found that selecting the top-ranked 30% (respectively 50%) of areas, a popular proposal for area-based conservation targets^6^, would conserve 62.4% (86.8%) of the estimated total carbon stock and 67.8% (90.7%) of all clean water provisioning, in addition to improving the conservation status for 69.7% (83.8%) of all species considered.

When optimizing conservation efforts for biodiversity only, we found that the groups that benefited the most were amphibian and plant species (Fig. 3b) and threatened species (Fig. 3c). The latter tend to have smaller range sizes and smaller absolute area targets than other groups and are inherently prioritized with area budgets ≤ 30% of land.

Our analysis included, for the first time in a global prioritisation analysis, a representative subset of plant distribution data totalling ~41% of described vascular plant species^32^ (Fig. 4). Incorporating data on plants resulted in spatial shifts in areas of importance for conservation, particularly in the western United States of America, West-Central and South Africa, South-West Australia, Central Brazil, as well as northern Europe and central Asian steppes and mountains compared to an analysis where plants are ignored (Fig. 4a). Overall we found montane and temperate grasslands, Mediterranean savannas and shrublands biomes to increase in importance when considering plants, whereas flooded grasslands and mangroves lost relative importance (Fig. 4b). The accumulation curves of species targets achieved were comparable between analysis variants with and without plants (Fig. 4c). Overall this indicates high surrogacy between vertebrate and plant species, despite spatial shifts in areas of importance (Fig. 4a).

**Fig. 4:**
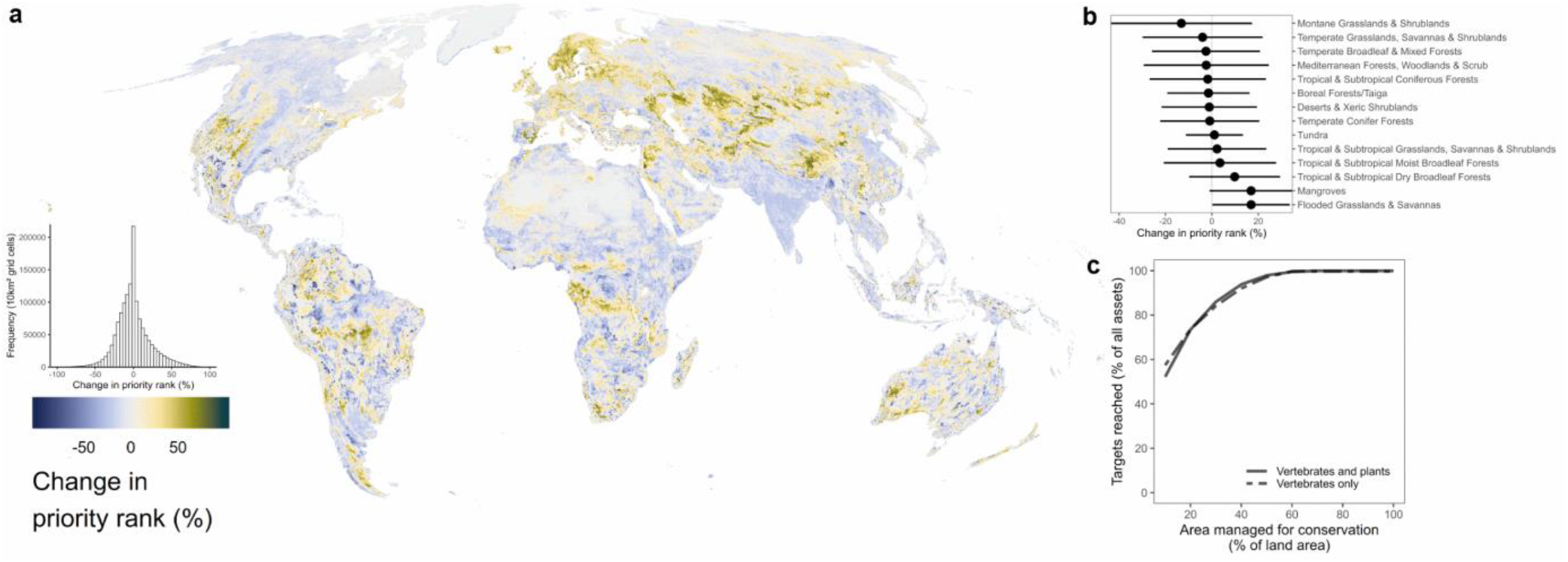
Change in global areas of biodiversity importance after adding plant species. (a) Calculated as the difference in areas of biodiversity importance with either plant species included or excluded. Positive changes (yellow to dark green) in rank imply an increase in priority if plant species are considered, while negative changes (light to dark blue) show a decrease in priority ranks. The map is at 10 km resolution in a Mollweide projection. (b) Average change in ranks per biome after plants have been added. (c) Representation curves of areas necessary to be managed for conservation with (solid) and without plants (dashed) included.

Areas of importance can vary spatially if species are given different weights, prioritising for instance the protection of threatened or more evolutionarily distinct species^20,21^. We tested the implication of prioritising the improvement of conservation status for these groups of species by weighting them by current conservation status or evolutionary distinctiveness. We found that doing so has only small inefficiency implications compared to a prioritisation without these weights (0.7% fewer biodiversity targets achieved when prioritising threatened species and 1.7% fewer when prioritising evolutionarily distinct species with 10% of land). Yet, overall spatial patterns of the top-ranked 10% of areas of importance were comparable, with only minor differences, notably highlighting the importance of New Zealand and the Brazilian Amazon for conserving threatened species, the Mediterranean Basin, North-West USA, Florida and fringes of the Amazon Basin for conserving evolutionarily distinct species (SI Fig. 10). These results highlight that threatened or more evolutionary distinct species are well covered by other species^30^, and their full conservation can be achieved at minimal extra cost.

## Discussion

How much area and where it should be managed for conservation is one of the key questions underpinning global biodiversity conventions and conservation planning discussions^4,29^. Our analyses suggest that even ambitious objectives such as ‘Half Earth’^24^ or ‘30 by 30’^6^ are insufficient to ensure that the conservation status of threatened species is improved and that non-threatened species remain so (Fig. 3). However, managing for conservation the top-ranked 30% of areas of importance for biodiversity, as identified here, can bring over 86% of the world’s terrestrial vertebrate and a representative sample of plant species (of ~41% of all plant species) to a non-threatened conservation status, with further increases in area offering minor additional returns (Fig. 3). Depending on the level of political ambition, an extra 20% of land could be dedicated to carbon storage as a contribution to climate regulation^6^ and sustainable management of natural resources. However, our analysis shows that considerable co-benefits can already be achieved by managing an optimally placed 30% of land, if conservation of biodiversity, carbon and water is planned for with spatial optimization approaches (Fig. 2). We caution that these estimates, and equally those from previous studies^6,14,16,23^, can vary with different data and methods applied.

We ranked priority areas in order of importance for conservation management; but we note that specific forms of management are highly contextual and will depend on local anthropogenic pressures, governance and opportunity costs. Areas of biodiversity importance that require strict protection and active management, e.g. where narrow-ranging and threatened species occur might be suitable for protected area expansion^31^. Other effective area-based conservation measures^32^, such as watersheds managed primarily for water resource management or community-managed forests, might be more suitable in areas where biodiversity, carbon and water benefits are high but threats to species conservation remain low.

Our analyses does not impose any constraint on feasibility or equity among countries^33^, some of which contain over half of their territory in the top-ranked 10% of global importance for biodiversity, carbon and water provision (Fig. 1). Thus, there is a need for fair resourcing of the required management actions to offset the financial burden on some, predominantly tropical, countries^33,34^. Existing funding mechanisms should further explore opportunities to synergistically benefit both biodiversity and NCPs, as has been shown in the case of carbon^26^. Future, synergistic conservation prioritization efforts should particularly focus on incorporating socio-economic constraints^35^, consider integrated scenarios of the projected distribution of biodiversity, carbon and water, support countries in identifying conservation actions at finer scale to maximize the achievement of national and global targets.

Our work also reveals research and data gaps in determining global areas of importance for terrestrial biodiversity conservation and NCPs. As NCPs we choose carbon and water because of their relevance to international conventions, but there are others we did not consider^22^ such as food provisioning or cultural relevance. Similarly, many aspects of biodiversity remain under-represented - although we consider a significant portion of plant species on Earth, and we developed a framework to remove spatial bias in priority setting resulting from incomplete taxonomic coverage - there is a need to expand available data on other groups such as freshwater, soil and invertebrate species^36,37^. We also only investigated the influence of evolutionary history on vertebrate, but not plant species, for whom hotspots of evolutionary history might differ, and ignored other dimensions such as functional rarity^38^. Despite remaining gaps in taxonomic coverage and species checklists, our analysis also confirms the results of previous, broad-scale studies^18,19,39^ that found high congruence between vertebrate and plant areas of importance, but we also highlight areas that would be overlooked if plants were not considered, especially so in dry grasslands, savannahs and Mediterranean shrublands (Fig. 4).

Our analyses highlight global areas of conservation importance that can maximize synergies across conventions (e.g. CBD, UNFCCC) and the SDGs. Particularly, our integrated maps could support governments in translating set targets (such as area-based conservation measures proposed for the 2021-2030 Strategic Plan of the CBD^4^) into national policies and actions on the ground and demonstrate how integrated spatial planning can be used to assist national biodiversity strategies. Meeting the SDGs requires real, transformative commitments that are yet to be enacted^1^, however, by maximizing synergies in efforts and resources, a pathway towards effective biodiversity conservation can be laid out for the next decade.

## Methods

### Biodiversity data

We utilized best available global species distribution data (overview in SI Table 1), including all extant terrestrial vertebrates and a representative proportion (41.31%) of all accepted plant species according to Plants of the World Online^40^. Extant mammal (5,685 species) and amphibian (6,660) distribution data were obtained from the International Union for Conservation of Nature Red List database (IUCN ver. 2019_2^41^), while bird (10,953) range maps were obtained from Birdlife International^42^. Data on the distribution of reptiles were obtained from the IUCN database when available (6,830 species), otherwise from the Global Assessment of Reptile Distributions (GARD) database (3,755^43^). We obtained native plant range maps (193,954 species) from a variety of sources, including IUCN, Botanic Gardens Conservation International (BGCI) and the Botanical Information and Ecology Network (BIEN). The IUCN and BGCI data contains expert-based range maps and alpha-hulls (see Supporting Information), while the BIEN data consists mainly of herbarium collections, ecological plots and surveys^44–52^, that were used to construct conservative estimates of species ranges using species distribution models (SDMs). We benefited from version 4.1 of BIEN, which includes data from RAINBIO^53^, TEAM^54^, The Royal Botanical Garden of Sydney, Australia, and NeoTropTree^55^. Additional plant plot data from a number of networks and datasets have been included in BIEN and a full listing of the herbaria data used can be found in the extended acknowledgements and online (http://bien.nceas.ucsb.edu/bien/data-contributors/all/). In cases where multiple data sources were available for the same plant species, we preferentially used expert-based range maps to characterize a species’ spatial distribution. A full description of the preparation and processing of the plant data can be found in the Supporting Information.

All vertebrate range maps were pre-processed following common practice^56^ by selecting only those parts of a species’ range where 1) it is extant or possibly extinct, 2) where it is native or reintroduced and 3) where the species is seasonally resident, breeding, non-breeding, migratory or where the seasonal occurrence is uncertain. We acknowledge that these ranges can contain some areas where the species is possibly extinct.

#### Suitable habitat refinement

Where data on species habitat and elevational preferences were available, we refined each species’ range to obtain the area of habitat (AOH) in which the species could potentially persist^57,58^. Data on species habitat preferences and suitable elevational range were obtained from the IUCN Red List database^41^ and, for an additional 1,452 reptile species in the GARD database, habitat preferences were compiled from an extensive literature search. For seasonally migrating birds and mammal species we ensured that separate habitat refinements were conducted for permanent and seasonally occupied areas of their range, that is, the breeding and non-breeding range. Whenever no habitat or elevation preferences were available for a given species, we used the full range except for areas considered to be artificial habitat type classes, such as arable or pasture land, plantations and built-up areas, noting that this could exclude areas suitable for some generalist species. For the AOH refinement we used a newly-developed global map (see Supporting Information) that follows the IUCN habitat classification system, thereby avoiding crosswalks between habitat preferences and land cover maps^59^. This data product integrates the best available land cover and climate data, while also using newly developed land-use data such as data on global forest management^60^. Finally, for each species and grid cell, we calculated the fractional amount (> 0-100%) of suitable habitat to include in the prioritisation analysis. Development of the habitat type map and all AOH refinement was performed on Google Earth Engine^61^.

#### Global representativeness

There is considerable bias and variability in the completeness of biodiversity records globally, particularly so for plant species^62^. To estimate the amount of geographic bias in completeness of distribution data among plants, we first estimated the proportion of species for which we had distribution data relative to the number of species known to occur in the regional checklists of World Checklist of Vascular Plants database^40^, which provides for each accepted species name its native regions from the World Geographical Scheme for Recording Plant Distributions (WGSRPD,^64^). We used geographic delineations for 50 WGSRPD level 2 regions^64^, but excluded Antarctica and mid-Atlantic islands (Saint Helena and Ascension) for which we had no plant records. The proportion of species for which we had range data varied from 11% in islands of the North pacific up to 100% in the Russian far east (mean 60.1% ∓ 24.5 SD). However, for 48 of the 50 WCSP regions we had distribution data for over >10% of all described plants known to occur natively in that region, (the exception being islands in the South-West and South-Central Pacific). For 44 of these 50 regions we had distribution data for >40% of described plants in those regions.

Having identified 10% as the minimum common denominator of completeness across most regions, we then used an iterative heuristic algorithm, to construct ‘representative’ subsets consisting of random samples that approximated 10% of species from each WGSRPD level 2 region while accounting for the fact that some species occur across multiple regions. To test if this approach yielded sets representative of biogeographic patterns of the full dataset, we compared the spatial patterns of scaled vertebrate species richness to the 10% sets of these species for each WGSRPD level 2 regions, random subsets of 10% of all vertebrates and for all vertebrates combined. We performed the test on vertebrates because we had range maps for ~95% of terrestrial vertebrates described, therefore we can assess if our subsampling to representative sets can replicate “true” patterns in species richness obtained with a complete sample of species in a taxonomic group. Spatial patterns of scaled species richness were identical across those sets, suggesting that this sampling approach can account for incomplete coverage (SI Fig 7a).

We also checked if the frequency distribution of range sizes within our subsets matched the range size distribution of the entire set using mammals as a test group, and found very modest differences between the full set and multiple subsets (SI Fig 7b). Having confirmed that this procedure recreates correct patterns of conservation priorities and it does not alter the range-size distribution (SI Fig 7), we proceeded to create 10 subsets of ~10% of plant species known to occur in each WGSRPD level 2 region and ten non-overlapping subsets of 10% of vertebrate species for all of our analyses. We found little difference among representation curves regardless of whether multiple representative subsets or all species were included in the SCP, although there was greater efficiency in the latter (SI Fig. 8).

### Carbon data

We used spatial estimates of the density of aboveground and belowground biomass carbon and vulnerable soil carbon^9^. Estimates for aboveground carbon (AGC) were created by selecting the best available carbon maps for different types of vegetation classes, identified spatially using the Copernicus Land Cover map in 2015^65^. We used Santoro *et al.* as a baseline for a global carbon biomass map^66,67^, which has been shown to be the most accurate, especially so for ‘tree’ covered land. In addition, we used more detailed estimates of above-ground biomass for African “open forest” and “shrubland” land cover^68^, global “herbaceous vegetation” and “moss and lichen” land cover^69^ and “cropland” and “bare/sparse vegetation” land-cover classes^70^. To map below-ground carbon, we applied corrected root-to-shoot ratios^71^ obtained from the Intergovernmental Panel on Climate Change (IPCC) technical guidance documents^72^. A newly developed forest management layer^60^ was used to update biomass density, by averaging estimates from 2010 and 2017^66^ in the most dynamic tree-covered classes (e.g. short rotation plantations, agroforestry).

The map of vulnerable soil organic carbon was created following IPCC Guidelines for National Greenhouse Inventories to estimate emissions and removals associated with changes in land use^72^. Vulnerable soil organic carbon was defined as those carbon stocks that could potentially be lost during the coming 30 years as a result of land use. We used recently published data on baseline soil organic carbon stocks^73^, and vulnerable stocks were estimated separately for mineral and organic soils. Organic soils were defined as those soils with ≥ 5% probability of being a Histosols according to USDA soil orders taxonomy^74^. All other soils were considered to be mineral soils. A 30cm depth was used to estimate vulnerable carbon stocks on mineral soils, while 200cm depth was used for organic soils. IPCC change factors (mineral soils) and emission factors (for organic soils) were used to estimate vulnerable soil organic carbon stocks according to IPCC land cover categories and climate zones. To be consistent with biomass carbon estimations, we created a crosswalk between the Copernicus global land cover map^65^ and IPCC land cover classes. The newly developed forest management layer^60^ was used to refine vulnerable carbon stock estimates for mineral soils, whilst managed forest with organic soils were excluded from this assessment given that due to drainage, these areas would be more suitable for restoration than for conservation action. Finally, all global carbon estimates were reprojected, summed and aggregated (arithmetic mean) to 10 km to match the biodiversity data in scale.

### Water data

For capturing water provisioning, we used estimates of potential clean water provision calculated by WaterWorld^75^ and Co$ting Nature^76^. This quantity calculates for each grid cell the volume of water available, as the accumulated water balance from upstream based on rainfall, fog and snowmelt sources minus actual evapotranspiration. Second, clean water was assessed using the Human Footprint on Water Quality (HFWQ) index, which is a measure of the extent to which water runoff is drawn from contaminating human land uses: both point (urban, roads, mining, oil and gas) and nonpoint (unprotected cropland, unprotected pasture) sources. The HFWQ index is calculated by cumulating the downstream runoff from polluting and non-polluting land uses and expressing the former runoff as a proportion of the total runoff. This is calculated by assigning an associated percentage (or dilution) intensity fraction to each land-use class (default values taken from^76^). The potential clean water provisioning service is calculated for each cell as the inverse of clean water (i.e. 100 - HFWQ) available from upstream. For the analysis we ranked each grid per river basin^77^ to determine their relative importance in delivering clean water within the basin.

### Prioritisation analysis

We determined global areas of importance to be managed for conserving biodiversity, carbon and water by using a spatial conservation prioritisation approach (SCP^78^). We divided the world in 10 km resolution ‘planning units’ (PUs, the cells of the land-surface area grids), in which ‘features’ are distributed (each species, plus carbon stocks and water provision), for which we establish conservation targets^79^. Each PU had an area ‘cost’ subject to ‘budget’ constraints (the total amount of the terrestrial land-surface within a PU). For biodiversity, we defined species-specific targets aimed at conserving the area of habitat (AOH) for a species to improve in conservation status (^15^, see Supporting Information) and for each species we calculated the amount of suitable habitat within each PU. For tonnes of carbon storage 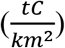 and/or volume of water 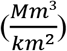, we maximized the total amount present in each PU. All PUs had a cost equivalent to the amount of land within them ({0 < *c* ≤ 1}), which we calculated from Copernicus land-cover data^65^. As global budget (B) we set different percentages of the terrestrial land surface area starting at 10%, then increasing by 10% increments up until all targets were met.

#### Problem formulation

Areas of importance for the conservation of biodiversity, carbon and water were determined by solving a global optimization problem. For each feature *j* included in the analysis we aimed to minimize the proportional shortfall^80^ in achieving each representation target *t_j_* given a planning unit cost *c* and an area budget *B* (10, 20, …, 100% of 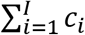 the planet). For all species, *t* is the target shortfall, that is, the difference between the part of an AOH that is included in the solution, and the amount that is necessary to be conserved for the species to improve in conservation status (^15^, Supporting Information), while for carbon storage and water provisioning *t* is the total amount available on the terrestrial land (the target is 100%). The problem is formulated as follows:

*Minimize* 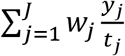

Subject to

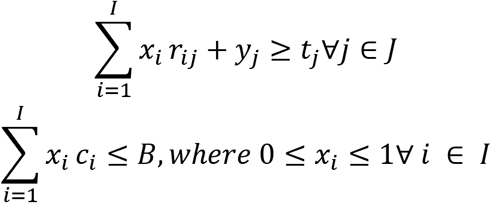

where *r_i,j_* is the amount (suitable habitat in km^2^, total tons of carborn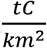or volume of water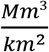) of feature *j* in planning unit *i, y_j_* is the shortfall for feature *j, t_j_* is the target for feature *j, c_i_* is the cost of grid cell *i* (the fractional area within the planning unit), *B* is the budget of the problem, *x_i_* is a proportional decision variable [0-1], where 1 means that the full PU and values ≥ 0 a fraction of the PU is selected, and W_j_ is the weight assigned to feature *j*. We tested different W_j_ of carbon, respectively water, relative to biodiversity and different weights among species based on their global threat status and/or evolutionary distinctiveness (Supporting Information). The problem is then solved for each budget incrementally, by ‘locking in’ previous solutions with lower area-budget prior to running the next prioritisation, effectively building nested sets of priorities with increasing budget *B*.

#### Analysis variants

For a separate analysis, we constrained the optimization by locking in the fraction of currently protected areas and adjusted the starting budget accordingly (Supporting Information). We then jointly optimized globally for biodiversity, carbon and water by minimizing the proportional shortfall^80^ in reaching the targets for each given area budget *B* (10, 20, …, 100% of the planet).

We furthermore considered a number of optimization variants in which we modified either the targets or weights assigned to each feature (biodiversity, carbon and/or water). For biodiversity, we also considered variants distinguishing between species intraspecific variation, threat status and evolutionary distinctiveness (SI Table 2). To capture intraspecific variation, we considered each part of a species range occurring in geographically separate biomes as a separate feature with its own target^28^, e.g. the Tiger (*Panthera tigris*) was split into five separate features, one for each of the five biomes overlapping the tiger range (Supporting Information). However, we only considered a split for features in which at least 2,200 km^2^ of AOH (the minimum absolute target area) was contained within a different biome compared to the biome with the majority of the species range. Compared to a version without these splits and when optimizing for biodiversity, carbon and water, overall differences were relatively minor (SI Fig. 11), but potentially locally important. We also collated data on species current threat status and, for vertebrates, data on their evolutionary distinctiveness (Supporting Information), and then calculated weights for each species following^13^. We then optimized all variants by minimizing the target-weighted shortfalls across all biodiversity features, subject to budget constraints.

We set weights for carbon storage and water provisioning relative to biodiversity in all analyses variants that included these assets. To do so we assigned sequences of weights from ‘none’ up to ‘equal’ importance by weighting carbon and water as follows: 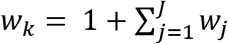, *w_k_* is the weight for carbon and water, *J* is the total number of species in the analysis, and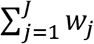 is the cumulative sum of all species weights. This weighting ensures that carbon is given equal importance to all species combined and that feature targets are treated equally in the optimization. We also created separate scenarios where *w_k_* is set to 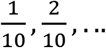 of the equal weighting relative to the cumulative shortfall for biodiversity. We visualized all scenarios with increasing budget and by the shortfall in carbon, water and improvement in species conservation status (Fig. 2) Because of the high computational cost of calculating (2*N_w_* − 1) * *N_B_* prioritizations, where *N_w_* is the number of weights and *N_B_* the number of budgets, for each of the 10 representative sets, we assessed differing weights at 50 km rather than 10 km resolution. However, we note that compared to a 10 km resolution, both spatial patterns and accumulation curves were highly similar (See Supporting Information and SI Fig. 9) and we don’t expect results to differ because of differences in resolution.

### Optimization algorithm and ranking

All SCP variants were solved using an integer linear programming (ILP) approach. Compared to other conservation planning solutions that rely on simulated annealing or heuristics^81^, ILP has been shown to outcompete those approaches in both speed and solution performance, being able to reliably find optimal solutions^82,83^. We ran all problem variants under each budgetary constraints (10%, 20%…100% of land), each with a representative set of species and solved them to optimality using proportional decisions (e.g. asking which fraction of a grid cell is part of the solution). For each problem variant, we therefore obtained 10 nested sets of priorities (priority ranks), each resulting from solving all budgetary constraints with a representative set of species. We summarized these priority ranks through an arithmetic mean while also separately calculating the coefficient of variation as a measure of uncertainty in priorities across representative subsets (SI Fig. 1). Selected planning units in the obtained solutions were investigated for the representation of input features by taxonomic group, threatened species and biomes.

All data preparation and analysis was conducted in R^84^ mainly relying on the ‘prioritizr’ package^85^ with the Gurobi solver enabled (ver 8.11,^86^).

## Data availability

All produced integrated maps will be made available through https://unbiodiversitylab.org/ and a data repository upon acceptance. The raw input data can be requested from the respective data providers, namely IUCN, GARD, Birdlife International, Kew Gardens and predicted plant distribution data will be made available as part of the BIEN initiative^44^. The IUCN habitat type map used to construct the AOH is made available in the Supporting Information. Any additional data not listed can be made available from the authors upon reasonable request or will be openly published separately.

## Code availability

Code to reproduce the main results will be made available upon acceptance.

## Acknowledgements

This work was conducted by the NatureMap consortium. We thank Richard Corlett & Tom Brooks who provided feedback on an earlier version of the manuscript. We furthermore thank Tom Hengl (OpenLandMap) for his advice on the Soil Organic Carbon analysis. This study has benefited from a number of data providers and networks. We explicitly acknowledge all data providers in a separate Extended Acknowledgements owing to their length (Supporting Information). The NatureMap project acknowledges funding from Norway’s International Climate and Forest Initiative (NICFI). Collection of the plant data used in this analysis has benefited from funding in form of a GEF grant 5810-SPARC ‘Spatial Planning for Area Conservation in Response to Climate Change’. CM acknowledges funding from NSF (National Science Foundation) grant DBI-1913673. RVG was supported by Australian Research Council DECRA Fellowship (DE170100208). Furthermore EAN and XF were funded by the Bridging Biodiversity and Conservation Science Program of the University of Arizona. NMH was supported by the European Union’s Horizon 2020 research and innovation program under the Marie Sklodowska-Curie grant agreement no. 746334. JCS considers this work a contribution to his VILLUM Investigator project “Biodiversity Dynamics in a Changing World” funded by VILLUM FONDEN (grant 16549) and his Independent Research Fund Denmark | Natural Sciences project TREECHANGE (grant 6108-00078B).

## Author contributions

MJ and PV designed the study, MJ led the analysis and interpretation of the data and has drafted the manuscript; PV conceived the study, contributed to the analysis and drafting of the manuscript; JH, BB,CM contributed to creating software used in the work; AA, CR, SGR, ML, DS, AvS, MM, JM, SP, IO, BS,CM,BJE,XF,PRR,BB,BM,RVG contributed to acquisition, analysis and interpretation of data; JH, MDM,JM,WJ,SR,JM,MO,MR,XDL contributed to interpretation of the data; GO, SM, ML, RG, MyL, OT contributed to acquisition and interpretation of data; XDL,VK, LM, NB, GW, JDS, GST contributed to conception of the study; VA,SPA,SCA,JRB,RTC,LH,PAM,JKM,DMN,NMH,EAN,DSP,PRR,JCS,CV,JJW provided data and contributed to interpretation of the data. All authors contributed to revising the manuscript. Correspondence and requests for materials should be addressed to MJ & PV.

## Supporting online material

### Material and Methods

#### Choice of resolution

We chose a spatial resolution of 10km to adequately capture global biodiversity and nature’s contribution to people per grid cell. For the biodiversity data we used estimates of a species global range. Previous studies have recommended coarser spatial resolution (~110km) when using species range maps as such, to better match equally downscaled atlas data considered to be the ‘true’ distribution of a species^1^, however, this can result in more costly prioritisations due to commission errors, without meaningful reductions in spatial biases^2^. In this study we refined a species range to an Area of Habitat (AOH,^3^) to minimize commission errors (false presences). This was done at a spatial resolution similar or even coarser than in comparable studies relying on the same range data^4–7^. Lastly, we also created separate maps of all analyses at 50km resolution to investigate differences on identified areas of biodiversity importance (SI Fig. 9), and found overall little to no difference between analyses done at these different resolutions. Nevertheless, we caution that the identified global areas of importance should not be used to inform conservation decisions on local or landscape scales.

#### Plant data preparation

To this date, there does not exist a single and consistent data source for species range data of all described plant species globally^8–10^. The total number of plant species globally is still unknown, with existing estimates ranging between 352,282 species^11^ and over 434,934 species^9^. To obtain a representative subset of described plant species, the NatureMap consortium gathered the best available plant distribution data from a variety of sources and types, acknowledging that none of them are without errors and biases, which we addressed by calculating spatially representative sets, each approximating the same proportion of species known to exist in an area, across the planet.

We first relied on expert-based global range estimates created by the International Union for Conservation of Nature (IUCN), Royal Botanic Gardens, Kew, and Botanic Gardens Conservation International (BGCI). For many plant species only curated point estimates of their range were available. Based on this data, range estimates were constructed using alpha-hulls, a generalization of convex hulls that are particularly useful for estimating species ranges whose habitat is irregularly shaped^12^ or where populations are spatially structured^13^. Parameters for alpha-hulls creation were adaptively selected, starting with initial alpha values - a parameter constraining the hull triangulation - of 2 or 3 recommended by the IUCN Red List categories and criteria, but adjusted for the distribution of records so that at least 95% of the records were included within the estimated range. The value of alpha ranges from zero (i.e. the finest resolution defined by the given set of points) to infinity (i.e. the coarsest resolution defined by the convex-hull). Since variations in alpha can also affect subpopulation structure (i.e. number of subpopulations), we combined alpha-hulls with the “1/10^th^ max” circular buffer method (i.e. the buffer size is set to the tenth of the maximum interpoint distance) to better capture subpopulation structure^13^. Finally, we limited the number of subpopulations to maximum of 10 and if the conditions above are not met (i.e. >= 95% of records inside the estimated range and <= 10 subpopulations), a minimum convex hull or a buffer built with the “1/10^th^ max” method is drawn around each record^13^. We split the occurrence records geographically into separate parts in cases the alpha hulls could not be constructed (for instance close to 180° longitude). In these cases, we applied the alpha-hull method to each individual dataset and merged the calculated hulls back into one unique range. All alpha hulls and “1/10^th^ max” buffers were created using the *rangeBuilder* package^14^. In total, data for 8,702 plant species ranges could be obtained through both sources, including 4,598 tree species from BGCI and 4,104 plant species from IUCN.

For plant species not yet assessed by IUCN or BGCI, we relied on modelled range estimates derived from occurrence records acquired through the Botanical Information and Ecology Network (BIEN) initiative, the Global Biodiversity Information Facility (GBIF.org 2019, https://doi.org/10.15468/dl.gvt20i) and from iNaturalist (www.inaturalist.org). Not all research grade observations from iNaturalist are transferred to GBIF and we thus downloaded all research grade iNaturalist plant data separately and merged them with the GBIF data, while removing duplicate observations.

The observations in the BIEN database are the product of contributions by 1,076 different data contributors, including numerous individual herbaria, and data indexers of herbaria (550+ are listed in Index Herbariorum), that were used to construct conservative estimates of species ranges using species distribution models (SDMs). For details of specimen data sources see^9,16^. We benefited from version 4.1 of BIEN, which includes data from RAINBIO^17^, TEAM^18^, The Royal Botanical Garden of Sydney, Australia, and NeoTropTree^19^. Additional plant plot data from a number of networks and datasets have been included in BIEN^8,9,16,20–25^ and a full listing of the herbaria data used can be found in the extended acknowledgements below and online (http://bien.nceas.ucsb.edu/bien/data-contributors/all/).

Taxon names associated with BIEN occurrence records were first corrected for misspellings, homonyms (e.g. plant and animal species with identical names) and synonyms. Afterwards all taxon names were standardized using TNRS v4.0 at default settings with checklists from Tropicos, The Plant List, USDA Plants, Global Compositae Checklist, ILDIS^26^. Standard BIEN preprocessing procedures furthermore ensure that species outside their native ranges were removed using lists of endemic taxa and the Native Species Resolver (NSR; https://github.com/ojalaquellueva/nsr). Observations were furthermore flagged and removed as cultivated based on keywords in the original observation metadata.

We applied the following preprocessing steps to all plant occurrence records from BIEN, GBIF and iNaturalist.We removed all occurrence records that (1) had no or impossible coordinates (e.g. < 90° S latitude or longitude >180° or <-180°), (2) had a coordinate uncertainty greater than 10 km, (3) had identical latitude or longitude coordinates, duplicate records or where coordinates had a precision smaller than one digit, (4) removed occurrence records in the vicinity (10 km distance) of country capitals or outside the lowest declared political division in the case of BIEN using the Geographic Name Resolution Service (GNRS; http://bien.nceas.ucsb.edu/bien/tools/gnrs/), near country or province centroids (1 km), or in the vicinity (1 km) of known zoos, botanical gardens or herbaria and (5) removed all occurrence points that fell into the open ocean^27^. For the modelling, we merged plant occurrence records from GBIF and iNaturalist into one dataset per species and only included those records from BIEN that were not already present in other data sources.

Plant species can have varying uncertainties in taxonomies and geographic spread and quite commonly occur in regions where the species is not considered native. In this study we relied on taxonomic and geographic information from the Plants of the World online (POWO) database, which provides for each accepted species name its native World Geographical Scheme for Recording Plant Distributions (WGSRPD) regions^28,29^. We only included plant species in the analysis whose name could be matched to POWO taxonomy (either as accepted name or as synonym) and which had at least one occupied grid cell in all WGSRPD level 2 regions in which the species is known to be native, to reduce influences of sampling biases. Lastly, we post-hoc removed from each predicted distribution all unconnected isolated patches outside native WGSRPD regions, which we identified through connected component labeling^30^.

For modelling plant species distributions we used a number of environmental covariates, which are adequate for the spatial scale (global at 10 km) of our modelling approach^31^. Data on present (1979-2013) climatic conditions (Annual Mean Temperature, Mean Diurnal Range, Annual precipitation, Precipitation seasonality, Precipitation of Warmest Quarter, Precipitation of Coldest Quarter, maximum accumulated Aridity (consecutive water deficit during months where potential evapotranspiration exceed precipitation) & estimated relative Precipitation of Warmest 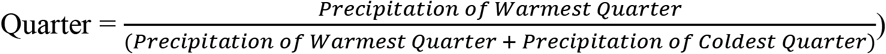 were obtained from CHELSA (http://chelsa-climate.org/,^32^). Data on global aridity^33^ and soil conditions (bulk density, % clay content, depth to bedrock, pH & % silt content all averaged over full depth to 200cm) from https://soilgrids.org^34^. These covariates were chosen based on their ecological relevance for plant species and on having global correlations < 0.7 with each other^35^. All environmental covariates were aggregated (arithmetic mean) to 10 km globally and projected to an equal-area Mollweide projection.

#### Point process modelling

For all plant species with 10 or more records available we fitted Poisson point process models (closely related to Maxent) using regularized down weighted Poisson regression models^36^, fitted with the R package glmnet^37^. We used up to a maximum of 20,000 background points in total, adjusted based on the total number of grid cells within the domain, and chose a spatial domain for predictions based on the biomes a species occurred in^38^. All candidate predictors were further filtered for collinearity for each individual species separately^35^, with highly collinear covariates (Pearson’ *r* > 0.7) within the domain removed.

Five independent folds were trained for cross validation, where folds were assigned based on spatial clusters to remove the influence of spatial autocorrelation on cross-validated performance statistics. Linear (all species), quadratic (species with >100 records), and product (species with >200 records) features were used. Regularization parameters for each model were determined based on one standard deviation below the minimum variance^37^. This resulted in five models per species which were then combined in an unweighted ensemble by calculating the arithmetic mean and standard deviation of the folds. Finally, the continuous predictions were thresholded to obtain binary presence/absence predictions based on the 5th percentile of the ensemble predictions.

#### Range-bagging models

For all plant species with between five and lower than ten records we utilized a ‘range bagging’ approach, which is a stochastic, hull-based method that can estimate climate niches from an ensemble of underfit models^39,40^, and is therefore well suited for smaller datasets. We randomly sampled 100 times a proportion *p* of records (*p* = 0.33, based on recommendations in^39^) and a subset *d* of environmental variables (*d* = 2,^39^). A convex hull is then projected around the subsampled records in environmental space, with a record considered part of the species range if its environmental conditions fall within the hull. We then chose a voting threshold of 0.165 (=0.33/2), implying that the grid cell is part of the species range at least half the time for each subsample. Upon visual inspection we generally found that this threshold leads to relatively conservative predictions. All range bagging records and environmental predictors were subjected to the same selection rules as for the point process models discussed above.

#### Grid cell data

For plant species with less than three covered grid cells records we used only those grid cells the points fall, which often describe the full distribution of the species known to science, many of which are globally rare^9^.

### Ancillary data

To account for current areas managed for conservation, we included data on current global protected areas from the global World Database on Protected Areas (WDPA, April 2019 version, IUCN and UNEP-WCMC 2019). Following commonly used WDPA preparation standards^41^, we excluded protected areas whose status was ‘proposed’ or ‘not reported’ and furthermore removed UNESCO Man & Biosphere reserves. This figure, however, does not include data from countries that have restricted the sharing of their dataset through the WDPA, such as China, Estonia, Saint Helena, Ascension and Tristan da Cunha^41^. All layers were first rasterized at 1 km, then aggregated to 10 km by calculating the relative fraction of area protected, so that small PAs were not lost. As a result, ~15% of the land surface was identified as being protected in the prioritisation analysis. Lastly, we prepared data on terrestrial biomes and ecoregions (http://ecoregions2017.appspot.com,^38^), which were likewise rasterized to 10 km resolution using a modal aggregation.

#### Habitat types map

Not all parts of a species range are equally suitable to allow a species to persist, thus requiring a refinement to an area of suitable habitat (AOH,^3,5^). In the past this refinement has commonly been attempted using a crosswalk^42^ between land-cover legends and habitat type information from the IUCN habitat type classification^43^. Crosswalks between different thematic legends can potentially cause issues such as inseparability of habitat types that are identical in land cover but different in climatic and soil conditions (e.g. tropical moist lowland forest and tropical mangrove forest). We developed a new global habitat type layer that follows the IUCN habitat classification system^43^. This layer is an intersection of the best currently available land cover dataset^44^, data on climate^45^ and other ancillary datasets, such as a novel data product on the distribution of global anthropogenically modified forests including tropical and temperate plantations (Lesiv et al. unpublished). Using this layer we refined all species ranges (see methods) at 1 km globally and calculated the fraction of suitable habitat per 10 km grid cell. We make a version of this global layer available as part of this manuscript^46^.

##### Prioritisation analysis

###### Target setting

One of the most impactful decisions in spatial conservation planning frameworks is the definition of feature targets. In the past, many studies set targets for species representation according to rules^47–49^ or area-based policies (e.g. 30% of a species range), which run the risk of leading to an excess of area for wide-ranging species and arbitrariness. We set targets relative to the amount of habitat necessary to improve a species conservation status as inspired by IUCN criteria^50^. We recognise that this only takes the range (area of suitable habitat) into account, and ignores other factors of extinction risk, such as population size and trends, but the purpose is to provide ecologically credible area-based conservation targets, rather than estimating extinction risk. For all species, these targets were defined as

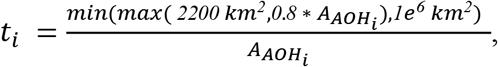

where *t_i_* is the relative target for a given species *i* and *A_AOH_i__*. the total area of suitable habitat for the species^50^. Whenever the numerator exceeded the *A_AOH_i__*. (e.g. is smaller than 2200 km^2^), the target was set to the whole AOH (100%), following^37^. In the prioritisation analysis we ranked each PU after formulating and solving a budget limited formulation of the reserve selection problem that aims to maximize conservation benefits.

###### Species-specific weights

Areas of biodiversity importance can vary depending on whether greater weight is placed on evolutionarily distinct^51^ and/or threatened species^52^. For this analysis we obtained data on the evolutionary distinctiveness (ED) scores for amphibians (99.7% of all species considered), birds (100%), mammals (100%) and reptiles (71.9%) from the EDGE program (EDGE 2019 list,^53^). For plant species there does not yet exist a species-resolved phylogeny^54^ and further research is necessary to fill that gap. Whenever ED scores could not be matched to species names, we used the congeneric or family-wide ED average^55^. ED scores represent the amount of unique evolutionary history of a species^56,57^, thus placing greater weight on evolutionary older and most distinctive lineages in a phylogeny. For example, Cuba and Hispaniola have evolutionary significance because these were the only two species of *Solenodon* that exist; the only members of the mammal family *Solenodontidae* which diverged from all other mammals over 60 million years ago, thus representing a disproportionate amount of evolutionary history. Data on the threat category (TC) of species was obtained from IUCN and encoded as numerical weight. In addition, for plant species we used data from the ThreatSearch online database^58^. We followed Pouzols et al. (2014) and assigned a weight of 8 to Critically Endangered species (CR), 6 to Endangered (EN), 4 to Vulnerable (VU), 2 to Near Threatened (NT) and 1 to species of Least Concern (1). Plant species without a standardized IUCN threat category, but which are considered threatened according to BGCI, were assigned a weight of 6. Species without sufficient current TC information or that were Data Deficient (DD) were assigned a conservative score of 2, given that many Data Deficient species are likely threatened with extinction^59,60^, especially so for plant species^11^. We separately incorporated for each species either the evolutionary distinctiveness (ED) score or the threat category (TC) as weight in the prioritisation, using weight from TC weights^52^. In total, we included data on ED weights for 34,308 species, TC weights for 43,211 species and calculated separated problem variants where data for both (29,780 species) is available (SI Fig. 10).

##### Supplementary figures and tables

**SI Fig. 1:**
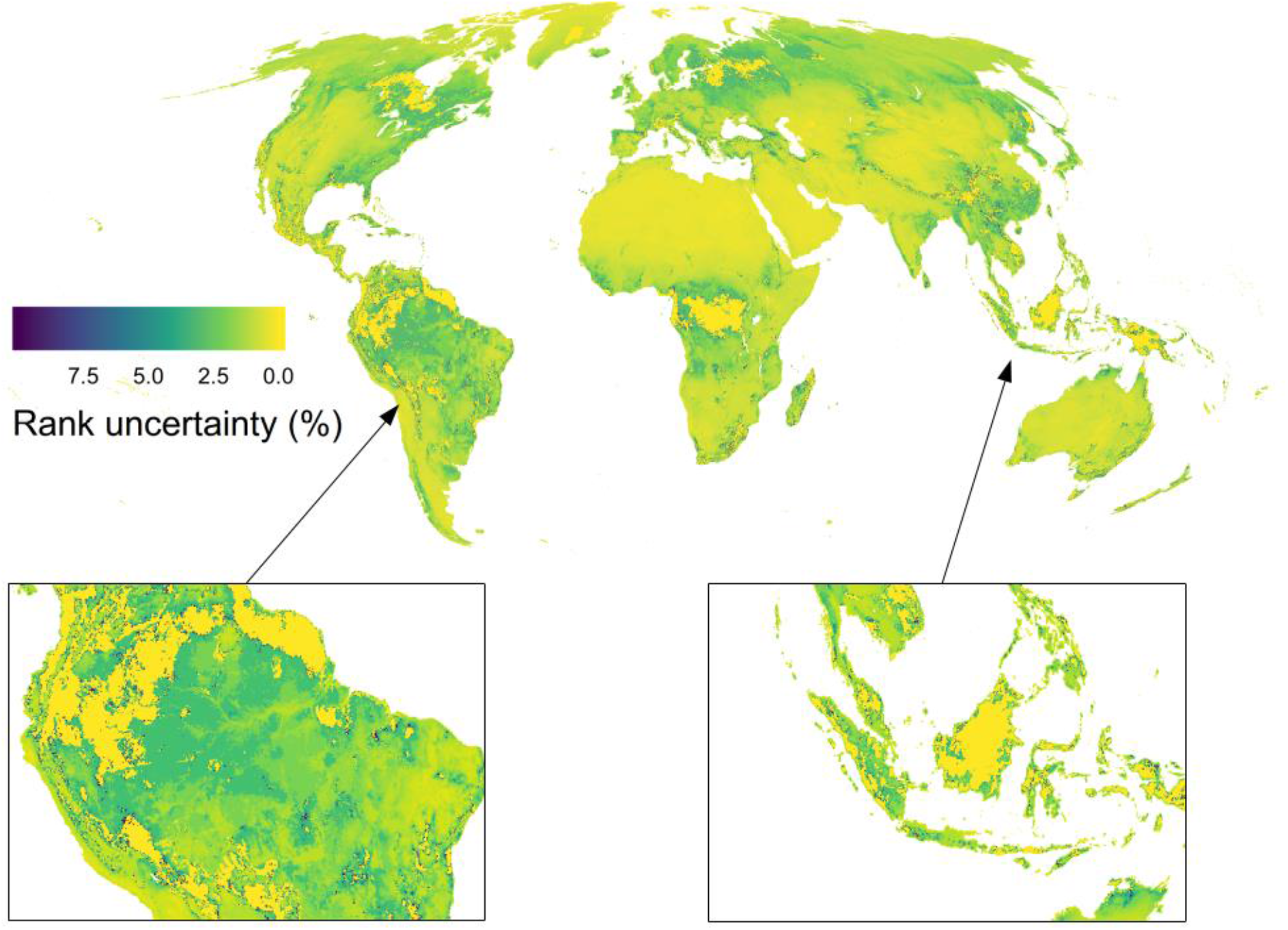
Uncertainty in ranks of areas of importance for biodiversity, carbon and water. Calculated as coefficient of variation across optimal solutions with different representative sets. Expressed as percentage with lower values indicating higher precision of ranks. Map can be interpreted as overall confidence in the mapped ranks (Fig. 1), given existing biases in species range data. Map is at 10 km resolution in Mollweide projection.

**SI Fig. 2:**
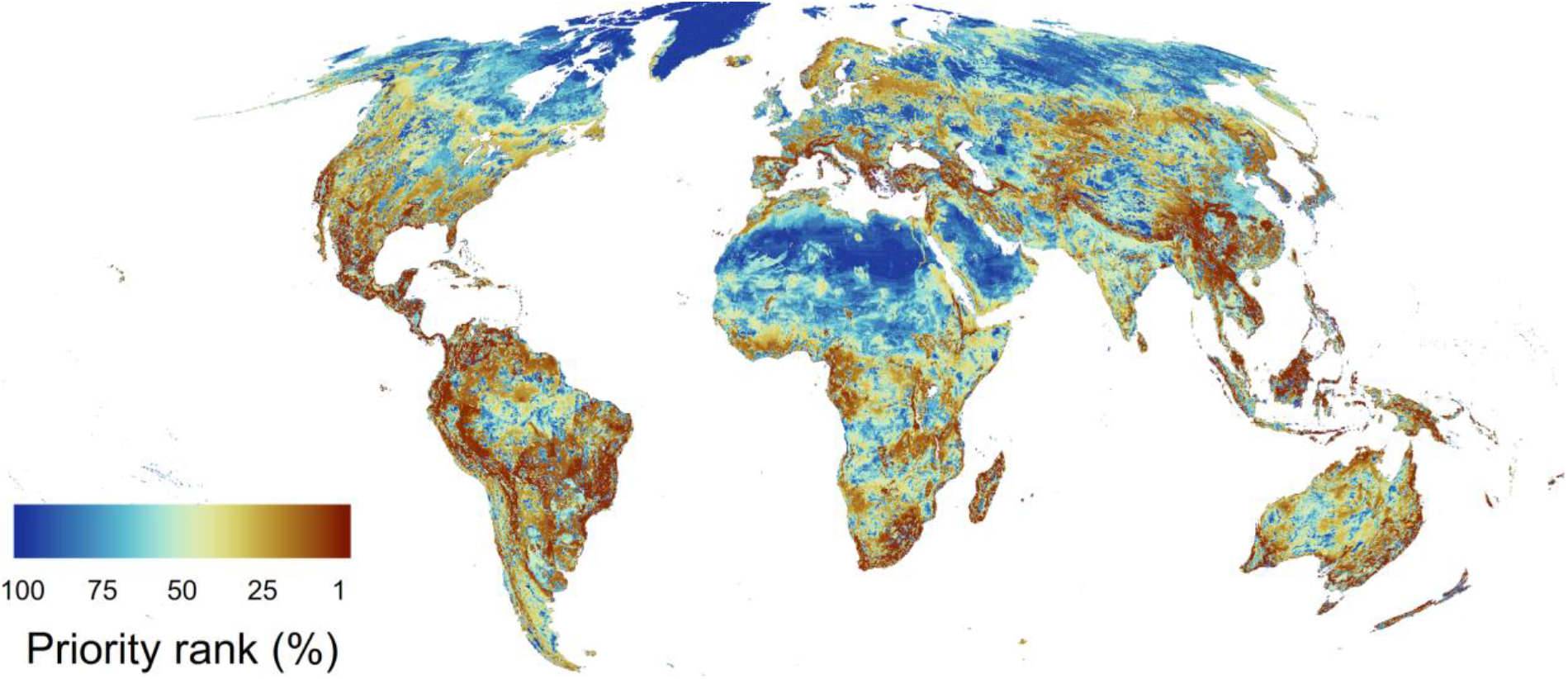
Global areas of importance for biodiversity only. Ranked hierarchical maps by the most (1-10%) and least important areas (90-100%) to conserve all of biodiversity globally. Map is at 10 km resolution in Mollweide projection.

**SI Fig. 3:**
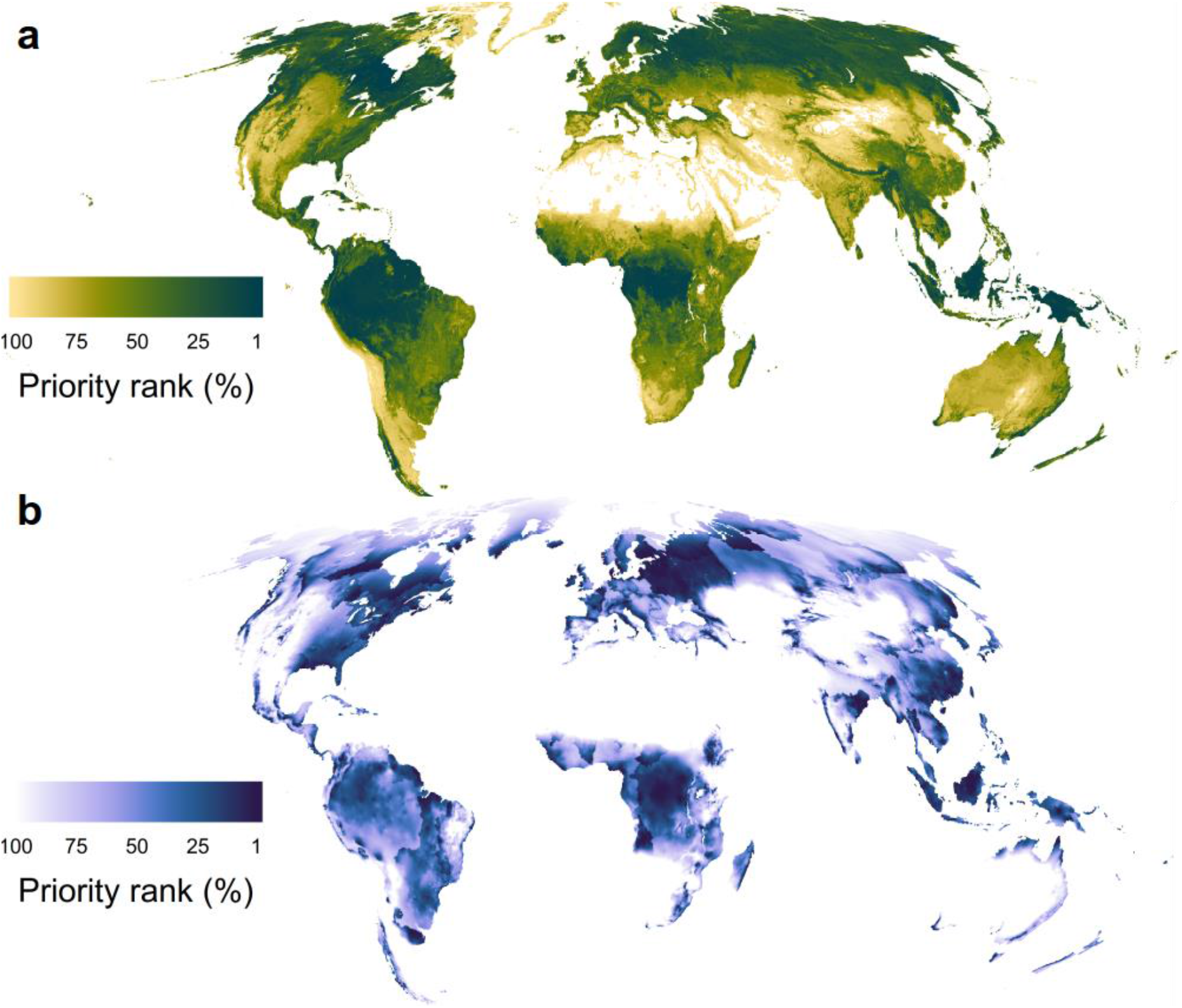
Global areas of importance for carbon and water. Normalized ranking for carbon (a) and water (b) presented as the most (1-10%) and least important areas (90-100%) to conserve globally. Map is at 10 km resolution in Mollweide projection.

**SI Fig. 4:**
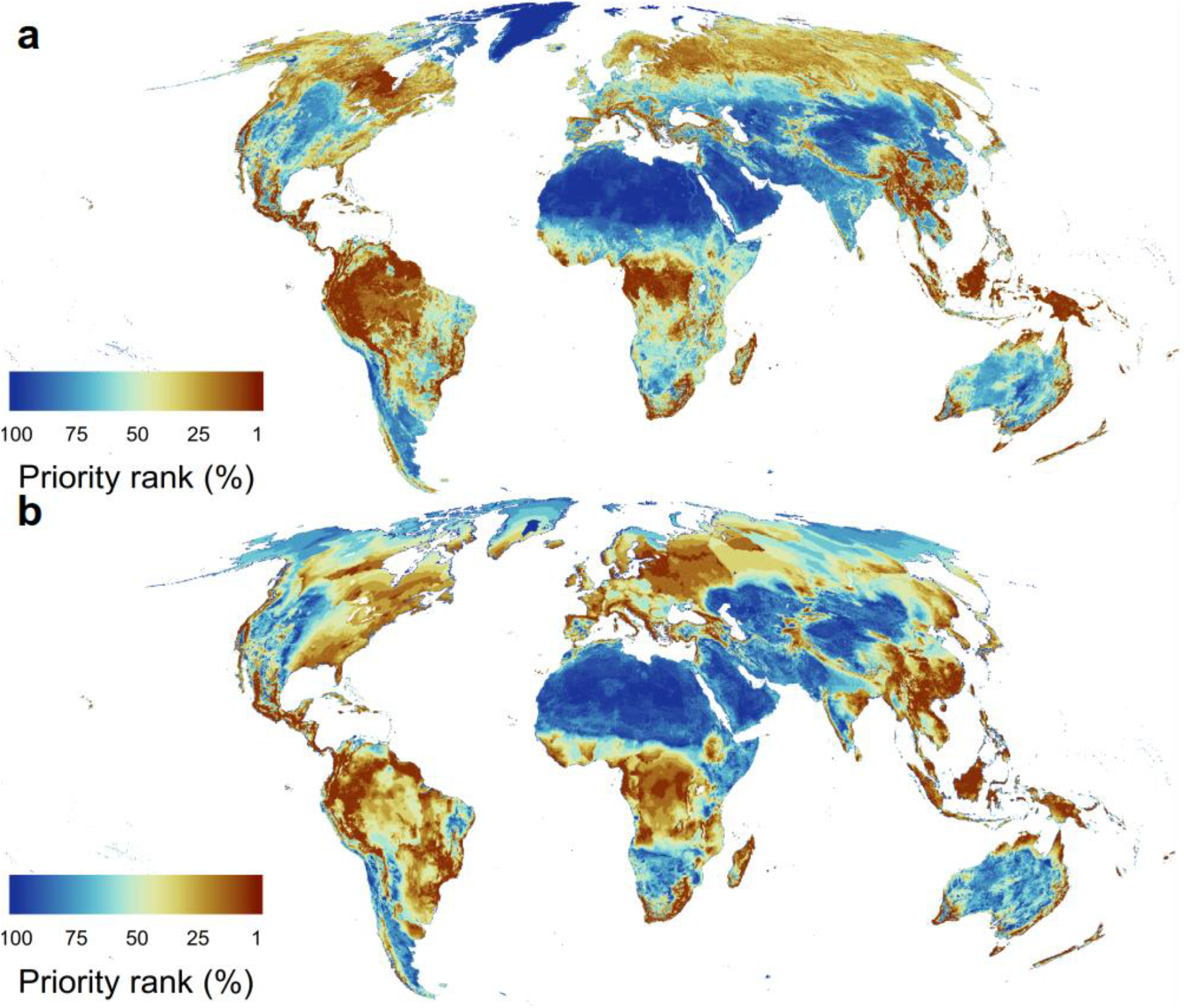
Global areas of importance for biodiversity and carbon or biodiversity and water. Showing an optimization across 10 representative sets for either (a) biodiversity and carbon or (b) biodiversity and water. All assets were jointly optimized and ranked hierarchical by the most (1-10%) and least important areas (90-100%) to conserve globally. Map is at 10 km resolution in Mollweide projection.

**SI Fig. 5:**
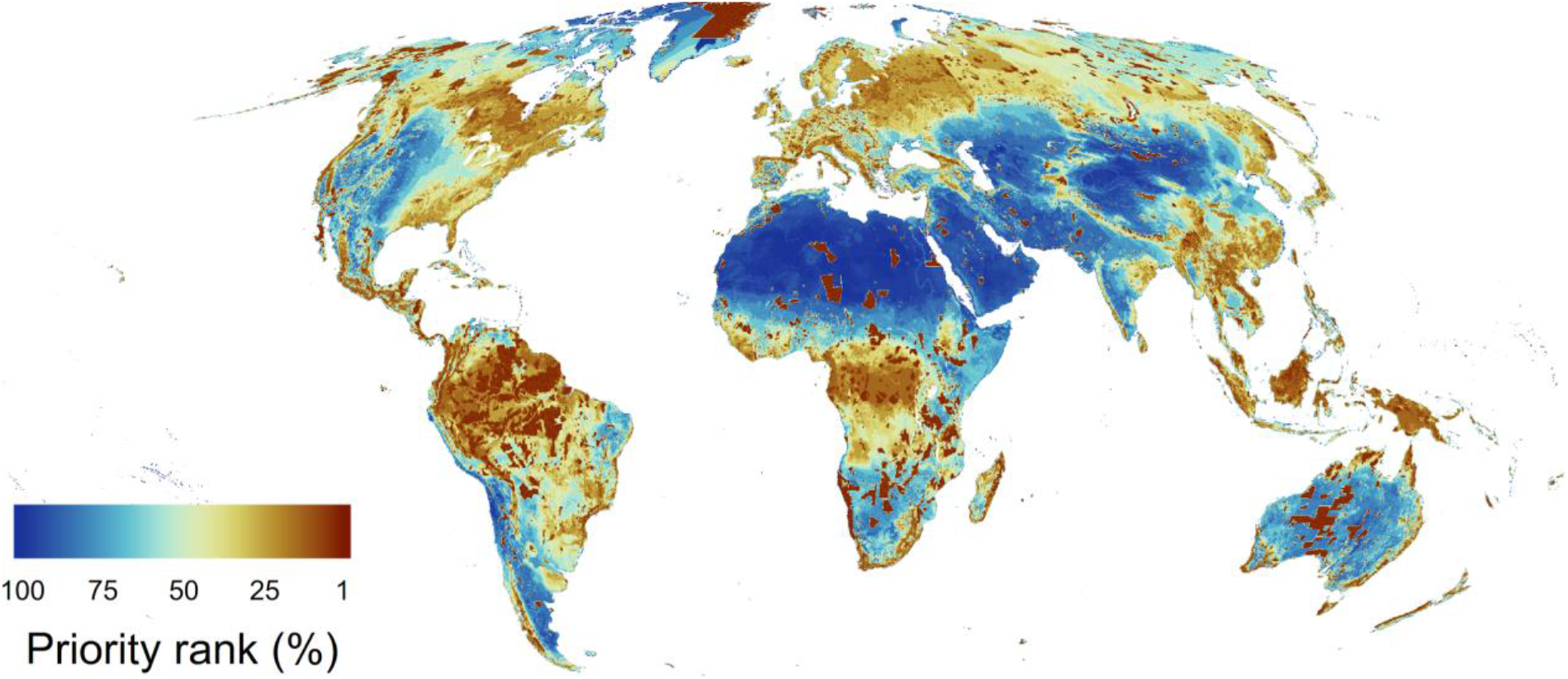
Global areas of importance for biodiversity, carbon and water considering current protected areas. All assets were jointly optimized and ranked hierarchical by the most (1-10%) and least important areas (90-100%) to conserve globally. The fraction of grid cells currently managed for conservation (https://www.protectedplanet.net) are considered to be part of the most important areas. Map is at 10 km resolution in Mollweide projection.

**SI Fig. 6:**
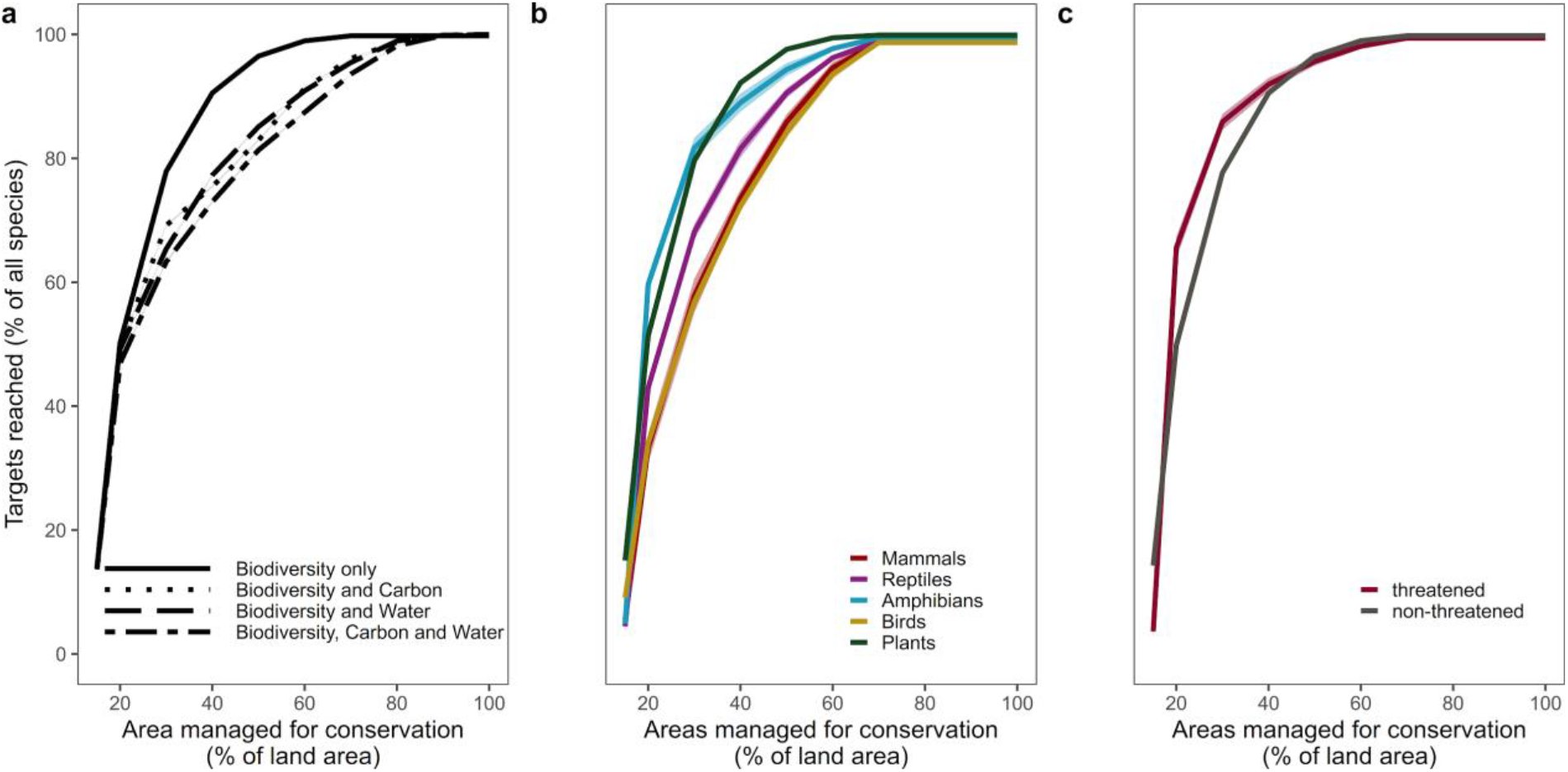
Accumulation curves showing how the number of species targets met increases with amount of land optimally allocated to conservation considering current protected areas. Shows the amount of land necessary for all assets to reach all persistence targets, defined as the amount of area needed for a species to be considered at reduced risk of extinction (see Methods). Uncertainty bands (~0.1% around the mean) show the standard deviation among representative sets. Estimates shown for species (a) overall and split by additional number of assets, (b) by taxonomic group, and (c) by current IUCN assessment of threat.

**SI Fig. 7:**
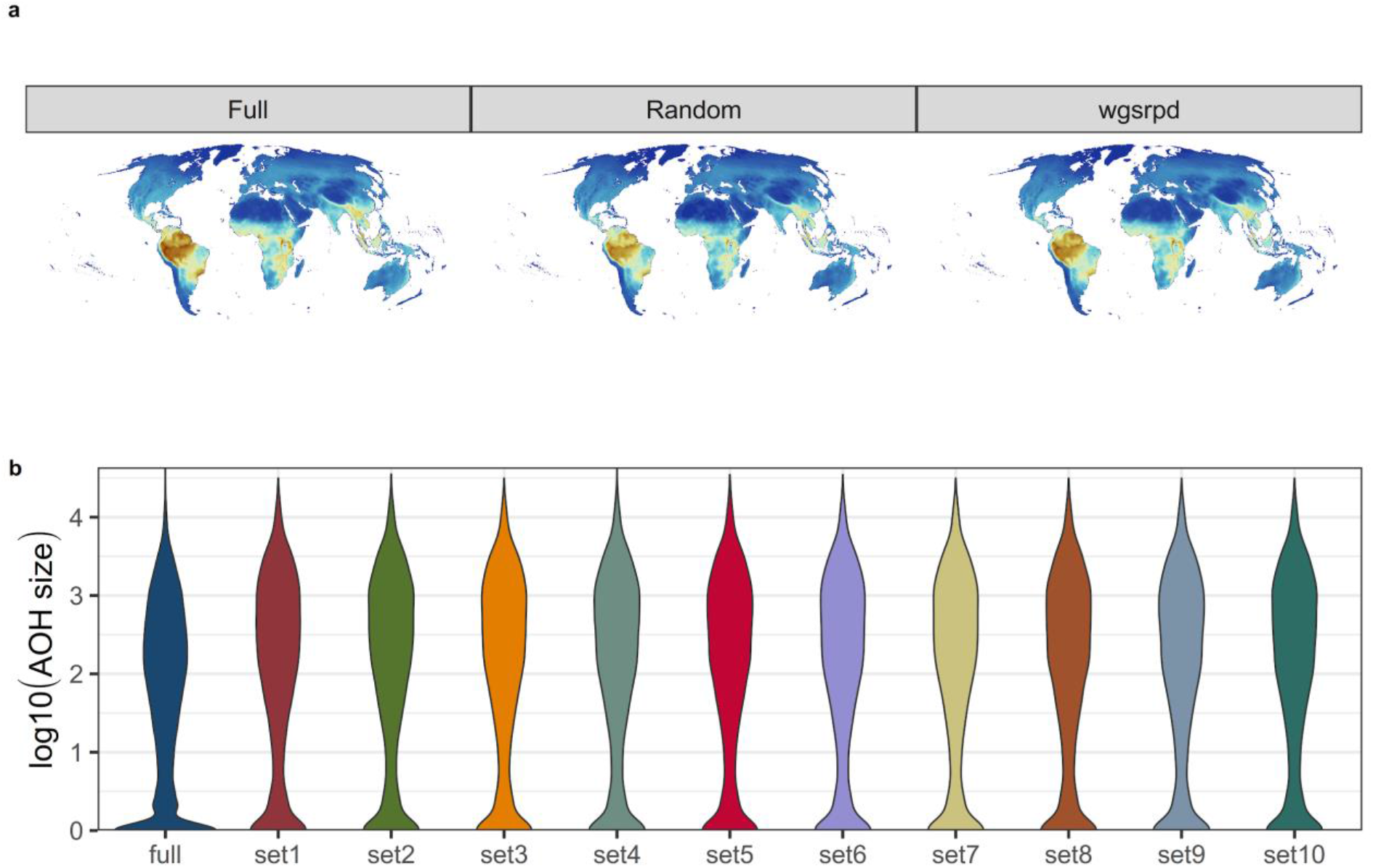
Comparison of representative sets spatially and in range size distributions. Compared to a full dataset, both subsampling at random and per WGSRPD region produces similar patterns in space and species area-size distributions. (a) Spatial map in Mollweide projection showing aggregated richness layers of all vertebrate species for the full dataset, a random sample and a representative sample by WGSRPD level 2 regions, (b) Shows the log10-transformed Area of Habitat (AOH) of all species in the full dataset (dark blue) compared to representative subsets of species (other colours).

**SI Fig. 8:**
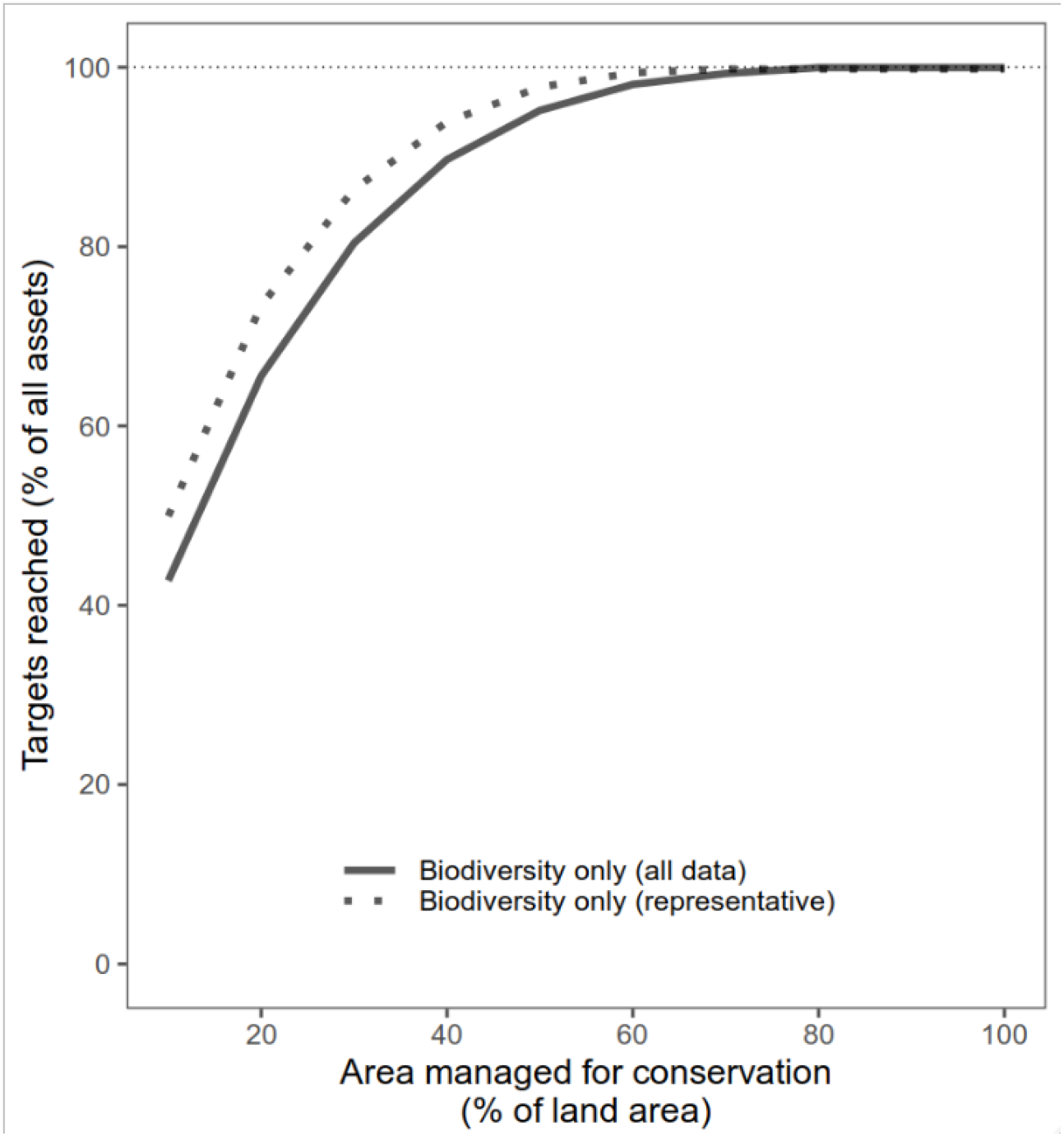
Accumulation curves showing how the number of species targets met increases with amount of land optimally allocated to conservation. Estimates shown for representative subsets (dotted line) and for all species included (solid line).

**SI Fig. 9:**
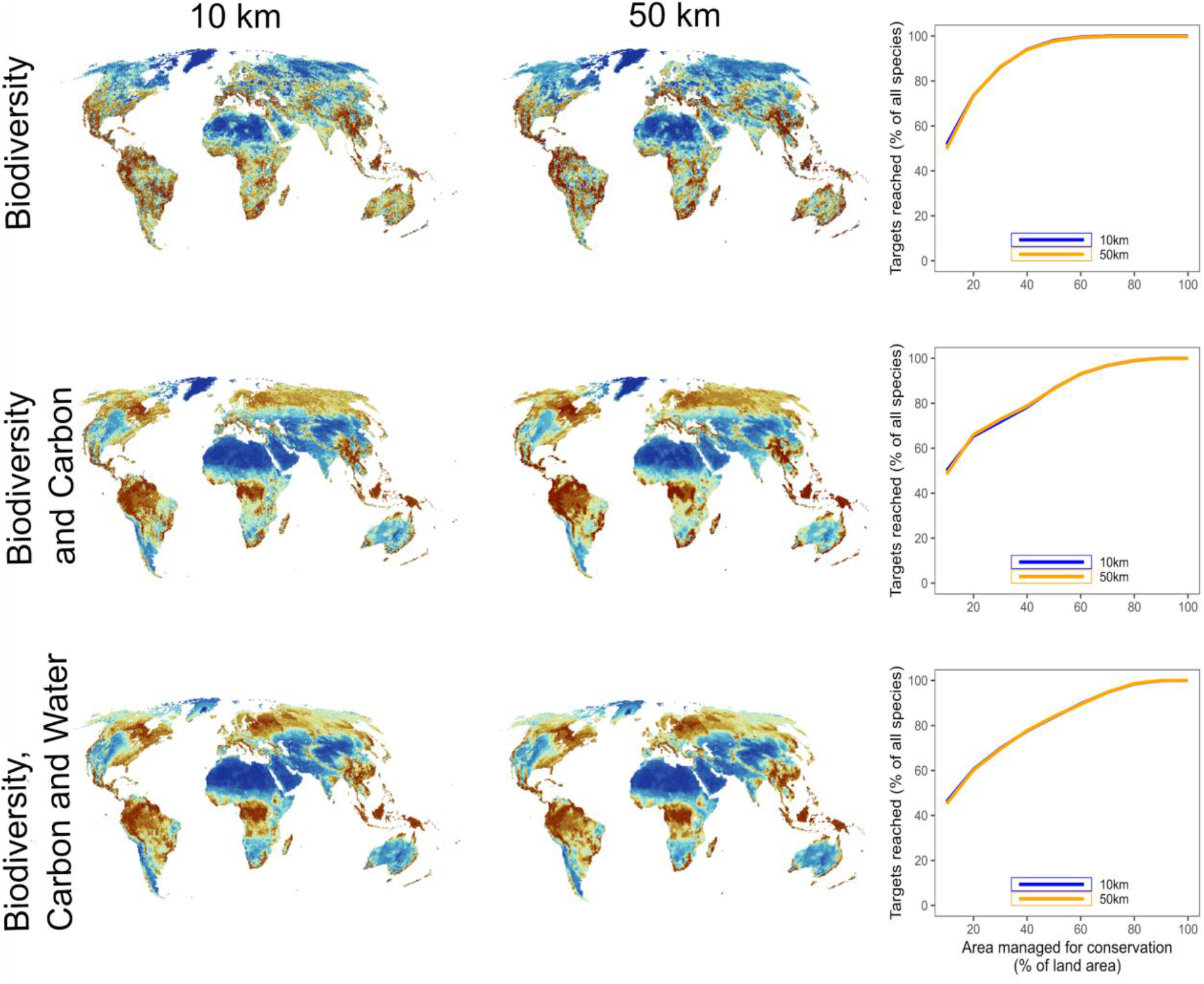
Comparison of global areas of importance at 10 km and 50 km areas. Comparisons in variants of areas of importance for biodiversity only; biodiversity and carbon; and biodiversity, carbon and water. Inset graphs show how the number of species targets met increases with amount of land optimally allocated to conservation for both 10 km (blue) and 50 km (orange).

**SI Fig. 10:**
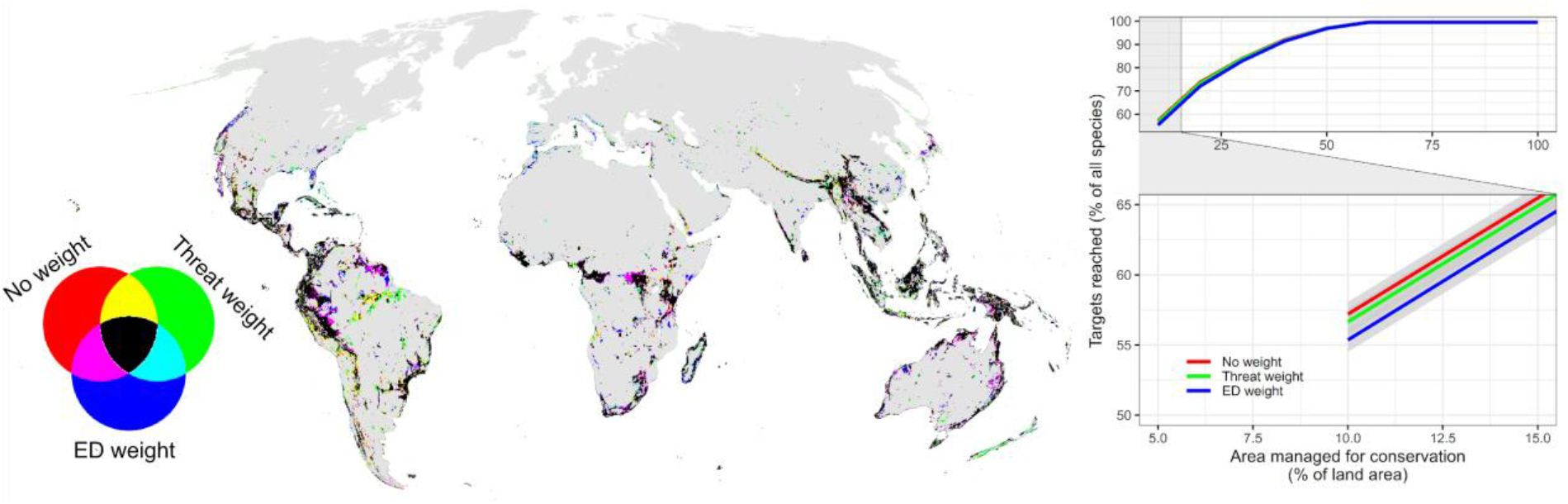
Difference in the top-ranked 10% solution for varying species weights. For each biodiversity feature a weight was assigned equating to either no differential weight (red), current threat category (green) or evolutionary distinctiveness (ED) (blue). Comparison was made only for species with data on both threat category and evolutionary distinctiveness. Grid cells coloured in black were selected in all three solutions. Map in Mollweide projection at 10 km resolution. The line plot shows the amount of land necessary for all species to reach all persistence targets, defined as the amount of area needed for a species to improve in conservation status (see Methods). Shown for either no weight (red), species weighted by threat status (green) and weighted by evolutionary distinctiveness (blue). The inset zoom highlights the difference among solutions at a budget of 10% terrestrial land area. The confidence bounds of accumulation curves indicate the uncertainty among representative sets.

**SI Fig. 11:**
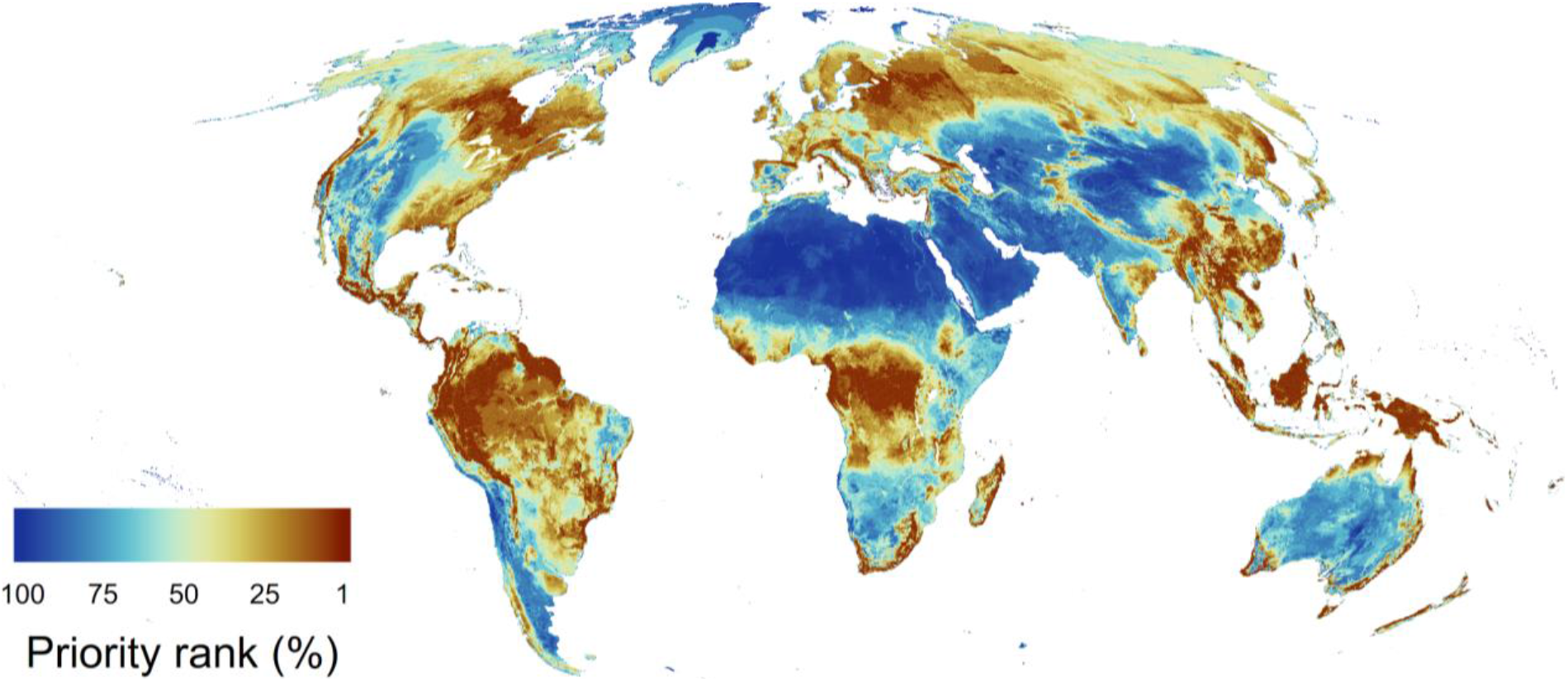
Global areas of importance for terrestrial biodiversity, carbon and water without biome splits. All assets were jointly optimized with equal weighting and ranked hierarchical by the most (1-10%) and least (90-100%) important areas to conserve globally. The map is at 10 km resolution in Mollweide projection.

**SI Table 1:**
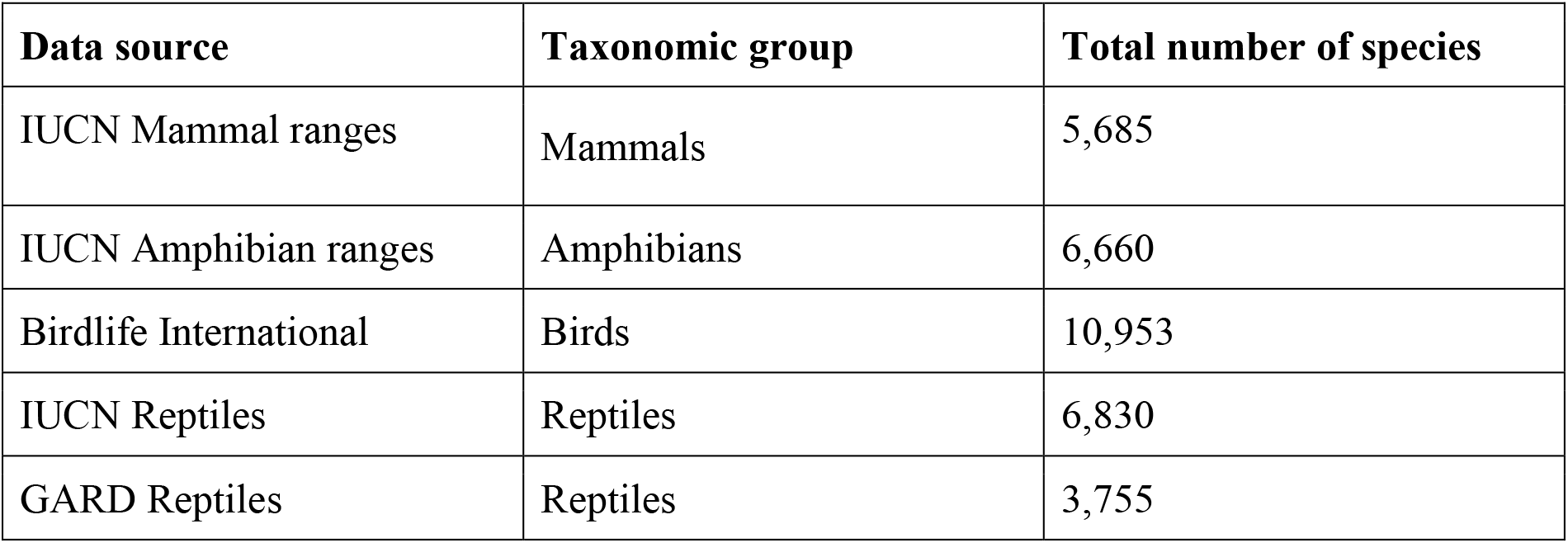
List of data sources included in the analysis. Shown is the source, taxonomic group and number of species ranges from that source. For the analysis we preferentially used species range data from IUCN and Birdlife International. Subsequently we relied on GARD, Kew and BGCI data and used BIEN estimates of species ranges for all other plant species not already included. Details on data preparation can be found in the methods and supporting information.

**SI Table 2:**
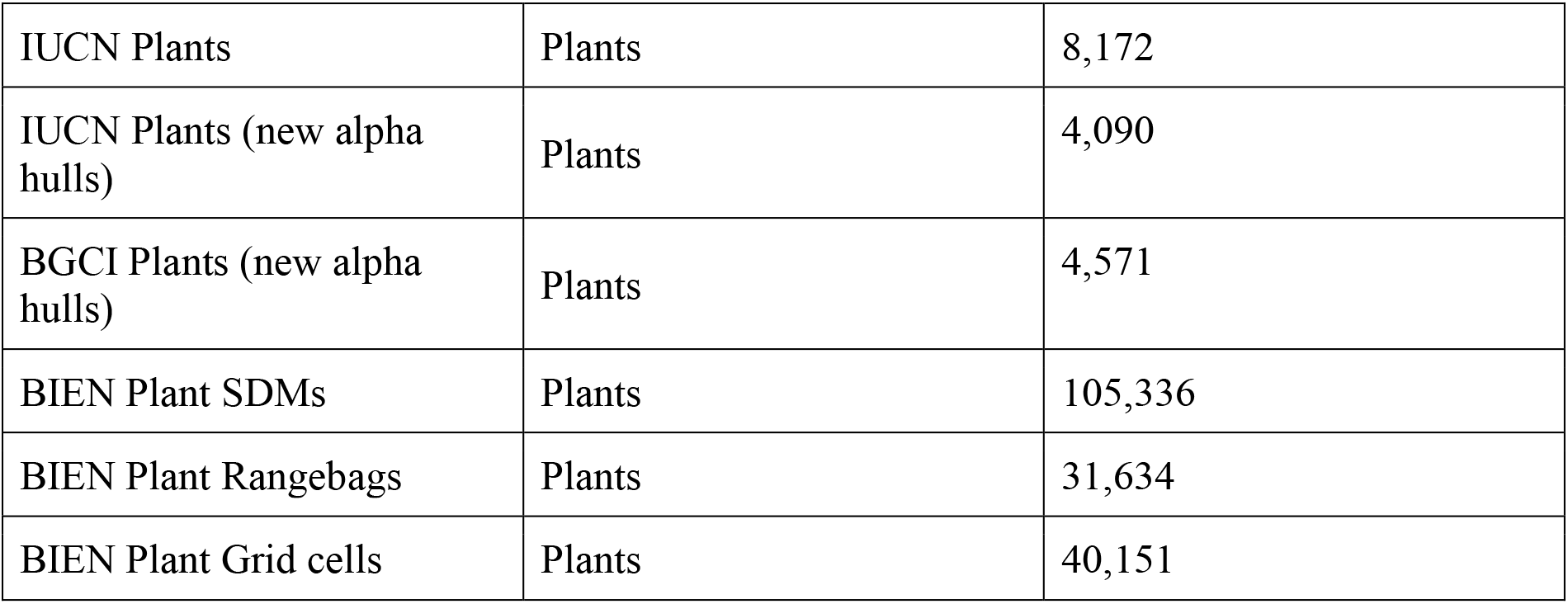
Problem variants created as part of the analyses. <uploaded separately>

## Extended acknowledgements

This study has benefited from data made available through a number of providers and networks. We would like to thank the IUCN Red List GIS Unit and Birdlife International for making vertebrate and plant species ranges available for scientific research. We thank the IUCN redlist and all species assessors globally for making habitat preference data available (https://iucnredlist.org). We thank Rikki Gumps and Claudia Gray for pointing us to the latest EDGE data (https://www.edgeofexistence.org/edge-lists/). We thank BGCI for making available up-to-date threat assessments of plant species via ‘threat search’ (https://tools.bgci.org/threat_search.php) and Royal Botanic Gardens, Kew for creating and making available the Global Plants of The World database (www.plantsoftheworldonline.org/).

We specifically thank Manaaki Whenua – Landcare Research and the New Zealand National Herbarium Network for making data available from the New Zealand National Vegetation Survey databank (NVS, https://nvs.landcareresearch.co.nz/) and the New Zealand Virtual Herbarium Network (NZVH). We thank iNaturalist for support in raising awareness and systematically collecting further data for countries with few plant occurrence records in a dedicated project (https://www.inaturalist.org/projects/naturemap-plants).

We thank all the 1076 data contributors to BIEN. This includes RAINBIO, TEAM, The Royal Botanical Garden of Sydney, Australia, and NeoTropTree. Plot data within BIEN are from the CVS, NVS, SALVIAS, VEGBANK, CTFS, FIA, MADIDI, and TEAM data networks and datasets (http://bien.nceas.ucsb.edu/bien/data-contributors/all). We acknowledge the herbaria that contributed data to BIEN: HA, FCO, DUKE, MFU, UNEX, VDB, ASDM, AMD, BPI, BRI, BRM, CLF, CNPO, L, LPB, AD, A, TAES, FEN, FHO, ANSM, ASU, B, BCMEX, RAS, RB, TRH, AAH, ACOR, UI, AK, CAS, ALCB, AKPM, EA, AAU, ALTA, ALU, AMES, AMNH, AMO, CHAPA, GH, ANGU, ANSP, ARAN, AS, CICY, BAI, CIMI, AUT, BA, BAA, BAB, CMMEX, BACP, BAF, BAJ, BAL, COCA, CODAGEM, BARC, BAS, BBS, BC, BCN, BCRU, BEREA, BG, BH, BIO, BISH, SEV, BLA, BM, BOCH, MJG, BOL, CVRD, BOLV, BONN, DAV, BOUM, BR, DES, BREM, BRLU, BSB, BUT, C, DS, CALI, CAN, CANB, CAY, EBUM, CBM, CEN, CEPEC, CESJ, CHR, ENCB, CIIDIR, CINC, CLEMS, F, COA, COAH,FCME,COFC,CP,COL,COLO,CONC,CORD,CPAP, CPUN, CR, CRAI, FURB, CU, G, CRP, CS, CSU, CTES, CTESN, CUZ, DAO, HB, DBN, DLF, DNA, DR, DUSS, E, HUA, EAC, EIF, EIU, GES, GI, GLM, GMNHJ, K, GOET, GUA, EMMA, HUAZ, ERA, ESA, FAA, FAU, FB, UVIC, FI, GZU, H, FLAS, FLOR, HCIB, FR, FTG, FUEL, GB, HNT, GDA, HPL, GENT, HUAA, HUJ, CGE, HAL, HAM, IAC, HAMAB, HAO, HAS, IB, HASU, HBG, IBUG, HBR, HEID, IEB, HIP, IBGE, ICEL, ICN, ILL, SF, HO, HRCB, HRP, HSS, HU, HUAL, HUEFS, HUEM, HUFU, HUSA, HUT, IAA, HXBH, HYO, IAN, ILLS, HAC, IPRN, IMSSM, FCQ, ABH, INEGI, INIF, BAFC, BBB, INPA, IPA, NAS, INB, INM, MW, EAN, IZTA, ISKW, ISC, ISL, GAT, JEPS, IBSC, UCSB, ISTC, ISU, IZAC, JACA, JBAG, JE, SD, JUA, JYV, KIEL, ECON, KSC, TOYA, MPN, USF, TALL, RELC, CATA, AQP, KMN, KMNH, KOELN, KOR, FRU, KPM, KSTC, LAGU, TRTE, KSU, UESC, GRA, IBK, KTU, ACAD, MISSA, KU, PSU, KYO, LA, LOMA, LW, SUU, UNITEC, TASH, NAC, UBC, IEA, GMDRC, LD, M, LE, LEB, LIL, LINN, AV, HUCP, QFA, LISE, MBML, NM, MT, FAUC, MACF, CATIE, LTB, LISI, LISU, MEXU, LL, LOJA, LP, LPAG, MGC, LPD, LPS, IRVC, MICH, JOTR, LSU, LBG, WOLL, LTR, MNHN, CDBI, LYJB, MOL, DBG, AWH, NH, HSC, LMS, MELU, NZFRI, MA, UU, MU, CSUSB, MAF, MAK, MB, KUN, MARY, MASS, MBK, MBM, UCSC, UCS, JBGP, DSM, OBI, BESA, LSUM, FULD, MCNS, ICESI, MEL, MEN, TUB, MERL, CGMS, MFA, FSU, MG, HIB, MIL, DPU, TRT, BABY, ETH, YAMA, SCFS, SACT, ER, JCT, JROH, SBBG, SAV, PDD, MIN, SJSU, MMMN, PAMP, MNHM, OS, SDSU, BOTU, OXF, P, MOR, POM, MPU, MPUC, MSB, MSC, CANU, SFV, RSA, CNS, WIN, MSUN, CIB, MUR, MTMG, VIT, MUB, MVFA, SLPM, MVFQ, PGM, MVJB, MVM, MY, PASA, N, UCMM, HGM, TAM, BOON, UFS, MARS, CMM, NA, NU, UADY, UAMIZ, UC, NE, NHM, NHMC, NHT, UFMA, NLH, UFRJ, UFRN, ULS, UMO, UNL, UNM, US, NMB, NMNL, USP, NMR, NMSU, WIS, NSPM, XAL, NSW, NT, ZMT, BRIT, MO, NCU, NY, TEX, U, UNCC, NUM, O, CHSC, LINC, CHAS, ODU, CDA, OSA, OSC, OSH, OULU, OWU, PACA, PAR, UPS, PE, PEL, SGO, PEUFR, PFC, PH, PKDC, SI, PLAT, PMA, PORT, PR, QM, PRC, TRA, PRE, PY, QCA, TROM, QCNE, QRS, UH, QUE, R, SAM, RBR, REG, RFA, RIOC, RM, RNG, RYU, S, SALA, SANT, SAPS, SASK, SBT, SEL, SIU, SJRP, SMDB, SMF, SNM, SOM, SP, SRFA, SPF, SPSF, SQF, STL, STU, SVG, TAI, TAIF, TAMU, TAN, TEF, TENN, TEPB, TFC, TI, TKPM, TNS, TO, TU, UAM, UB, UCR, UEC, UFG, UFMT, UFP, UGDA, UJAT, ULM, UME, UNA, UNB, UNR, UNSL, UPCB, UPEI, UPNA, USAS, USJ, USM, USNC, USZ, UT, UTC, UTEP, UWO, V, VAL, VALD, VEN, VMSL, VT, W, WAG, WAT, WII, WELT, WFU, WMNH, WS, WTU, WU, Z, ZSS, ZT, CUVC, LZ, AAS, AFS, BHCB, CHAM, FM, PERTH, SAN.

